# A neuronal relay mediates muscle-adipose communication that drives systemic metabolic adaptation to high-sugar diets

**DOI:** 10.1101/2023.08.15.553340

**Authors:** Olga Kubrak, Anne F. Joergensen, Takashi Koyama, Stanislav Nagy, Mette Lassen, Jacob Hald, Dennis Madsen, Kenneth V. Halberg, Michael J. Texada, Jakob L. Hansen, Kim Rewitz

## Abstract

Obesity leads to impaired insulin signaling and tissue sensitivity, which drive the onset of type 2 diabetes. Insulin resistance leads to a reduction in cellular glucose uptake, resulting in elevated blood glucose levels, which consequently cause β-cell dysfunction and development of diabetes. Although improving insulin signaling is a key target for restoring whole-body glucose homeostasis and reversing diabetes, the multi-organ mechanisms that regulate insulin signaling and tissue sensitivity are poorly defined. We screened the secretome and receptome in *Drosophila* to identify the underlying interorgan hormonal crosstalk affecting diet-induced insulin resistance and obesity. We identified complex interplay between muscle, neuronal, and fat tissues, mediated by the conserved BMP and LGR signaling pathways, which augments insulin signaling and improves dietary sugar tolerance. We found that muscle-derived BMP signaling is induced by sugar and governs neuronal Bursicon signaling. Acting through its LGR-family receptor, Bursicon both enhances insulin secretion and improves insulin sensitivity in adipose tissue, thereby preventing sugar-induced hyperglycemia. Inhibition of Bursicon-LGR signaling in adipose tissue exacerbates sugar-induced insulin resistance, and we discovered that this condition could be alleviated by suppressing NF-κB signaling. Our findings identify a muscle-neuronal-fat tissue axis that drives metabolic adaptation to high-sugar conditions by modulating insulin secretion and adipocyte insulin sensitivity, highlighting mechanisms that may be exploited for the development of strategies for the treatment and reversal of insulin resistance.

## Introduction

Insulin signaling is essential for glucose homeostasis. In peripheral tissues, insulin promotes the uptake of glucose from the blood and stimulates its conversion into glycogen and fat for storage. Insulin-stimulated glucose uptake and metabolism are reduced in insulin-resistant subjects, leading to elevated blood glucose levels^1^. This elevation intensifies the secretory demand on pancreatic β-cells, which is further amplified by the increase in body mass due to obesity. This can result in β-cell loss or damage, ultimately leading to inadequate levels of circulating insulin and the development of diabetes^2^. The adaptive mechanisms of hormonal and cellular physiology effectively manage glucose homeostasis under normal conditions. However, obesity and high-calorie diets rich in sugars can trigger insulin resistance in adipose tissue and other organs^3^. Despite the great medical importance of this condition, the underlying hormonal pathways and mechanisms that become disrupted due to nutrient excess and subsequently lead to impaired insulin signaling remain poorly defined.

Many *in-vivo* studies and models of diet-induced insulin resistance have linked decreased tissue insulin sensitivity with factors such as lipid accumulation, endoplasmic-reticulum stress, and inflammatory stress responses^4,5^. While improving β-cell function and ameliorating insulin resistance in obese conditions are key steps towards restoring normal glucose absorption to prevent and treat diabetes, the underlying mechanisms involve multi-organ crosstalk within complex hormonal systems that are still inadequately defined. Animal models of diet-induced obesity display insulin resistance and thus enable a range of experimental approaches to elucidate these mechanisms and identify targets for diabetes treatment^6,7^. Because of conservation of most metabolic pathways and many hormonal systems, the fruit fly *Drosophila* has become a valuable model for understanding the contribution of nutrition to metabolic disorders. In flies, diet-induced obesity produces many of the same pathophysiological effects observed in humans with obesity, indeed including hyperglycemia, altered insulin secretion, and insulin resistance^8,9^. Given the fly’s genetic adaptability, this model of diet-induced insulin resistance can be exploited to provide insight into the complex tissue crosstalk that drives insulin resistance and diabetes progression.

Insulin sensitivity is essential in both muscle and adipose tissue for glucose uptake and storage, and these tissues are critical regulators of systemic glucose homeostasis^10,11^. Signals from organs such as the gut and adipose tissue also relay nutritional information to the brain, which produces factors that regulate systemic energy homeostasis^12^. The crosstalk between these and other organs is mediated by circulating hormones and cytokines. Gut hormones such as glucagon-like peptide 1 (GLP-1) are responsible for incretin effects, which potentiate glucose-induced insulin release^13^. While gut-derived signals such as GLP-1 have become a primary therapeutic target for obesity and diabetes treatment, diverse myokines and adipokines also mediate crosstalk between the muscles, adipose tissue, and other organs, including the brain, to modulate systemic energy metabolism and insulin sensitivity^14,15^. The adipokine leptin can improve hyperglycemia and diabetes in animal models^16^, and myokines can enhance insulin sensitivity^17^. Like those of mammals, the tissues of the fly produce a range of adipokines, myokines, and gut-derived hormones that govern development, metabolism, and food choice^18–24^. Mammalian leptin is secreted from adipose tissue and stimulates insulin secretion and sensitivity in other organs via the nervous system. Similarly, the *Drosophila* leptin analog Unpaired 2 is secreted from the adipose tissue and promotes insulin secretion^23^. While the adipokine leptin enables fat tissue to modulate insulin signaling and sensitivity in the liver and muscles, reciprocal communication from muscles to adipose tissue via the nervous system has not been described.

Here, we present the results of an extensive *in-vivo* knockdown screen in *Drosophila* that examines the effects of the secretome and receptome on sucrose toxicity. Our findings show that, on a high-sugar diet that induces obesity-like phenotypes including insulin resistance, interorgan communication between muscle, neuronal, and fat tissue maintains insulin production and adipose insulin sensitivity, which mitigates sugar-induced hyperglycemia. In response to elevated sugar levels, muscle-derived Bone Morphogenetic Protein (BMP) signals to neurons expressing the hormone Bursicon. This factor acts through Leucine-rich-repeat-containing GPCR (LGR) signaling and directly stimulates insulin secretion and indirectly boosts insulin signaling by enhancing the sensitivity of adipose tissue to insulin. Moreover, adipose insulin sensitivity is inversely regulated by NF-κB signaling.

## Results

### *In-vivo* RNA-interference screen of the secretome and receptome for hormonal signals affecting sugar tolerance

The maintenance of sugar homeostasis in response to nutritional challenges requires the coordinated adaptive functions of multiple organs. Secreted molecules such as hormones and cytokines mediate signaling between organs to orchestrate these coordinated responses. To identify potential interorgan signaling factors that mediate the effects of nutrient excess on insulin signaling and sugar tolerance, we performed an *in-vivo* RNAi-mediated screen of the secretome and receptome in a fly model of diet-induced obesity and diabetes. In this system, a high-sugar diet (HSD) induces phenotypes that mirror some effects of human obesity, including hyperglycemia and insulin resistance^8,9^. The fly genome encodes only a single insulin receptor, and thus this pathway combines the effects of mammalian insulin itself (sugar balance) and IGFs (tissue growth). Because of this common signaling channel, insulin-perturbation effects in the larva manifest as a strong sugar-dose-dependent growth retardation that leads to developmental delay (Fig. 1a,b). We rationalized that knockdown of genes with critical roles in sugar tolerance would impair survival or delay pupariation on HSD (5x sugar in the diet) while having no effect on development on a normal diet (ND, 1x sugar). We confirmed that loss of Hexokinase-A (Hex-A), the enzyme mediating the first step in the glycolytic breakdown of glucose, produces a sugar-dose-dependent developmental-delay phenotype, validating our screening approach for identifying genes important for sugar metabolism (Fig. 1c). To ensure that our approach could detect differences in timing despite variation in the number of offspring among the different fly crosses, we assessed whether crowding would affect HSD-induced phenotypes. Examining mean pupariation timing in 25-mm diameter vials of 1x- or 5x-sugar medium containing between 5 and 150 larvae, we observed no difference in pupariation time between populations (Extended Fig. 1), indicating that larval crowding does not affect timing within this range, which covers the distribution of populations produced within our screen.

**Figure 1.**
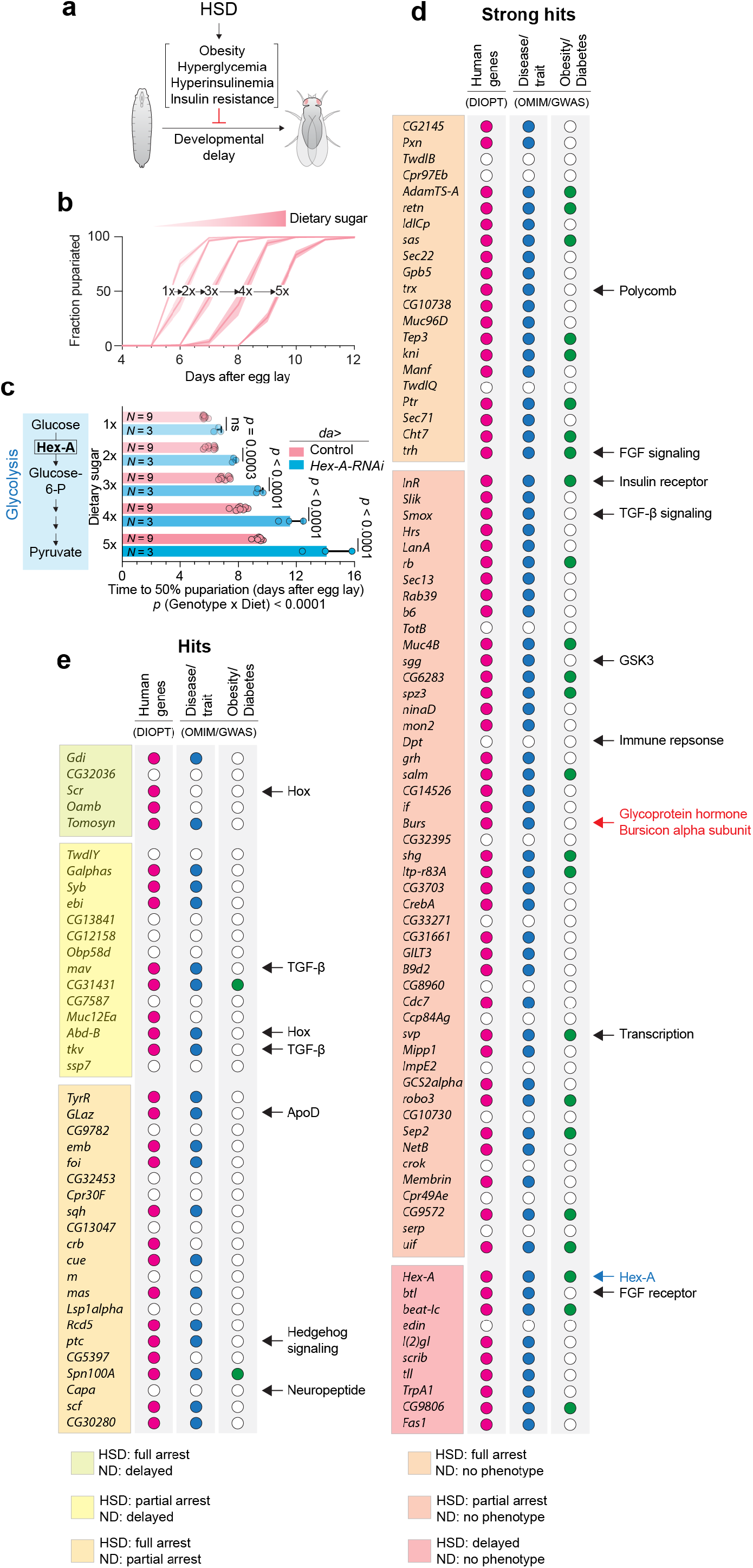
A screen for genes affecting adaptation to high-sugar diet. (A) The basis of the screen. (B) Pupariation timing on media containing a range of sugar concentrations. (C) Top: the time until 50% of animals had pupariated, in controls and *Hex-A* knockdown animals, on a range of dietary sugar levels. Bottom: the difference in 50%-pupariation times between controls and knockdowns on 1x (normal diet, ND) and 5x sugar (high sugar diet, HSD). (D) Screen hits that displayed no phenotype on 1x sugar while exhibiting full (top) or partial (middle) failure to pupate or a delay in pupariation (bottom) on 5x-sugar medium. Among these hits was the Burs (Burs-alpha) subunit of the Bursicon glycoprotein heterodimer. Other notable hits are indicated as well. Genes marked with a red dot have at least one ortholog in humans; if the human gene(s) have been associated with a disease or metabolic trait, the *Drosophila* gene is marked with a blue dot; and if that human disease or trait is related to obesity or diabetes, the *Drosophila* gene is marked with a green dot. (E) Screen hits that displayed a stronger phenotype on 5x sugar than on 1x sugar – those that permit development, albeit with delay, on 1x sugar while fully (top) or partially (middle) blocking development on 5x, and those that exhibit a partial arrest on normal diet and a full arrest on 5x sugar (bottom). Genes are flagged as in D, and notable hits are indicated. Statistics: Error bars represent mean and SEM. ns, not significant (*p*>0.05). C, top: two-way ANOVA with Šidak’s correction; bottom, one-way Kruskal-Wallis nonparametric ANOVA.

Next, we screened for genetically induced alterations in sugar-induced developmental delay – indicating a role in sugar tolerance or insulin signaling – by targeting 2,256 genes encoding potential secreted factors and receptors. Hits were identified by comparing the HSD-induced delay of knockdown animals to effects on development of controls. To identify sugar-dependent phenotypes, we compared 5x-sugar (HSD) phenotypes to 1x-sugar (ND) phenotypes. Global knockdown driven by *daughterless-GAL4* (*da>*) identified 119 genes (a hit rate of 5.3%) whose reduction exacerbated the sugar-induced delay phenotypes or impaired survival, or both, on HSD, compared to controls (*da*>+), but whose manipulation had no or comparatively minor effects on ND (Fig. 1d,e). The significant phenotypes ranged from full developmental arrest on HSD to partial arrest or enhancement of the HSD-induced developmental delay. Among the strongest 79 hits that produced only sugar-dependent phenotypes (*i.e.*, developmental arrest or enhanced delay on HSD paired with no delay on ND; Fig. 1d), 81% have at least one human ortholog (predicted by DIOPT^25^) (Fig. 1d). While many of these sugar-dependent gene hits (39%) have been associated with diabetes, obesity, or lipid and glucose metabolism (DIOPT-DIST based on OMIM and GWAS disease terms^25^), only 5% (2 genes) of the genes showing partially sugar-dependent phenotypes were associated with these terms (Fig. 1e). Among the sugar-dependent screen hits were several genes known to be important for sugar tolerance and metabolic homeostasis, including those encoding the insulin receptor (InR) itself; the transcription factor Seven-up (svp), a positive regulator of insulin signaling in adipose tissue required for maintaining sugar homeostasis under high-dietary-sugar conditions^26^; Smad on X (Smox), a TGF-β-pathway transcription factor required for glycemic control on a high-sugar diet via regulation of glucagon-like signaling in the fat tissue^27^; and FGF signaling components, which regulate insulin in response to fat-tissue oxygen levels^28^. These observations validate our approach and suggest that it is a powerful strategy to identify the mechanisms and hormonal pathways that control sugar homeostasis. The Hedgehog and Capa-peptide signaling pathways that have been linked with metabolism were also identified ^29,30^, along with other interesting pathways that have not previously been linked with sugar homeostasis, such as Hox genes (Scr and Abd-B), Polycomb chromatin remodeling, and immune responses. Among our strongest hits, we identified α-Bursicon, which with its paralog β-Bursicon [encoded by *Burs* and partner of bursicon (*Pburs*), respectively] makes up the heterodimeric cystine-knot glycoprotein Bursicon. This hormone acts through the conserved leucine-rich repeat-containing G-protein coupled receptor (Lgr) dLgr2/Rickets (orthologous with mammalian Lgr4/-5)^31^, as an important factor for sugar tolerance.

### Neuronal Bursicon signaling is required for metabolic adaptations that prevent dyslipidemia and hyperglycemia on diets rich in sugar

To investigate the role of Bursicon signaling in sugar tolerance, we first assessed the effects of a variety of *Burs* loss-of-function genotypes on the HSD phenotype. We found that animals homozygous for a *Burs* null mutation (*Burs^Z4410^*) affecting the entire animal exhibited a strong prolongation of the HSD-dependent delay, as did tissue-specific RNAi-mediated knockdown or CRISPR-mediated somatic deletion of *Burs* in Burs^+^ cells (driven by *Burs-GAL4*, *Burs*>) (Fig. 2a,b and Extended data Fig. 2a). Burs, the α-subunit of the heterodimeric Bursicon hormone, was previously shown to be expressed without the β-subunit in the adult midgut and to control systemic metabolism in response to sugar feeding^32^. However, during development, the Bursicon hormone is believed to function as a heterodimer produced by neurons^31^. Consistent with this, Burs and Pburs are predominantly expressed in the nervous system during development^33^. The α-subunit encoded by *Burs* is expressed at very low levels in the larval midgut, and the β-subunit encoded by *Pburs* is not detectably expressed. To address whether neuronal Burs:Pburs signaling is required for sugar tolerance, we disrupted *Burs* and *Pburs* specifically in the nervous system and assessed sugar-dependent effects. Animals with pan-neuronal knockdown or knockout of either *Burs* or *Pburs* showed a sugar-dependent developmental delay (Extended data Fig. 2b,c). These data suggest that it is the heterodimeric Burs:Pburs hormone, produced by neurons expressing both subunits, that is important for metabolic adaptations to high-sugar conditions. We further corroborated our findings by feeding animals a diet with a high sugar-to-yeast ratio but with normal sugar concentration and low yeast concentration (6% sugar and 0.34% yeast ratio, *i.e.*, 0.1x yeast, the latter being the primary source of protein and sterols). In comparison to the controls, animals with neuronal knockdown or knockout of *Burs* demonstrated a substantial delay on the low-yeast diet (Extended data Fig. 2d). This further supports to the importance of neuronal Bursicon signaling for adaptation to diets with a high ratio of sugar to yeast.

**Figure 2.**
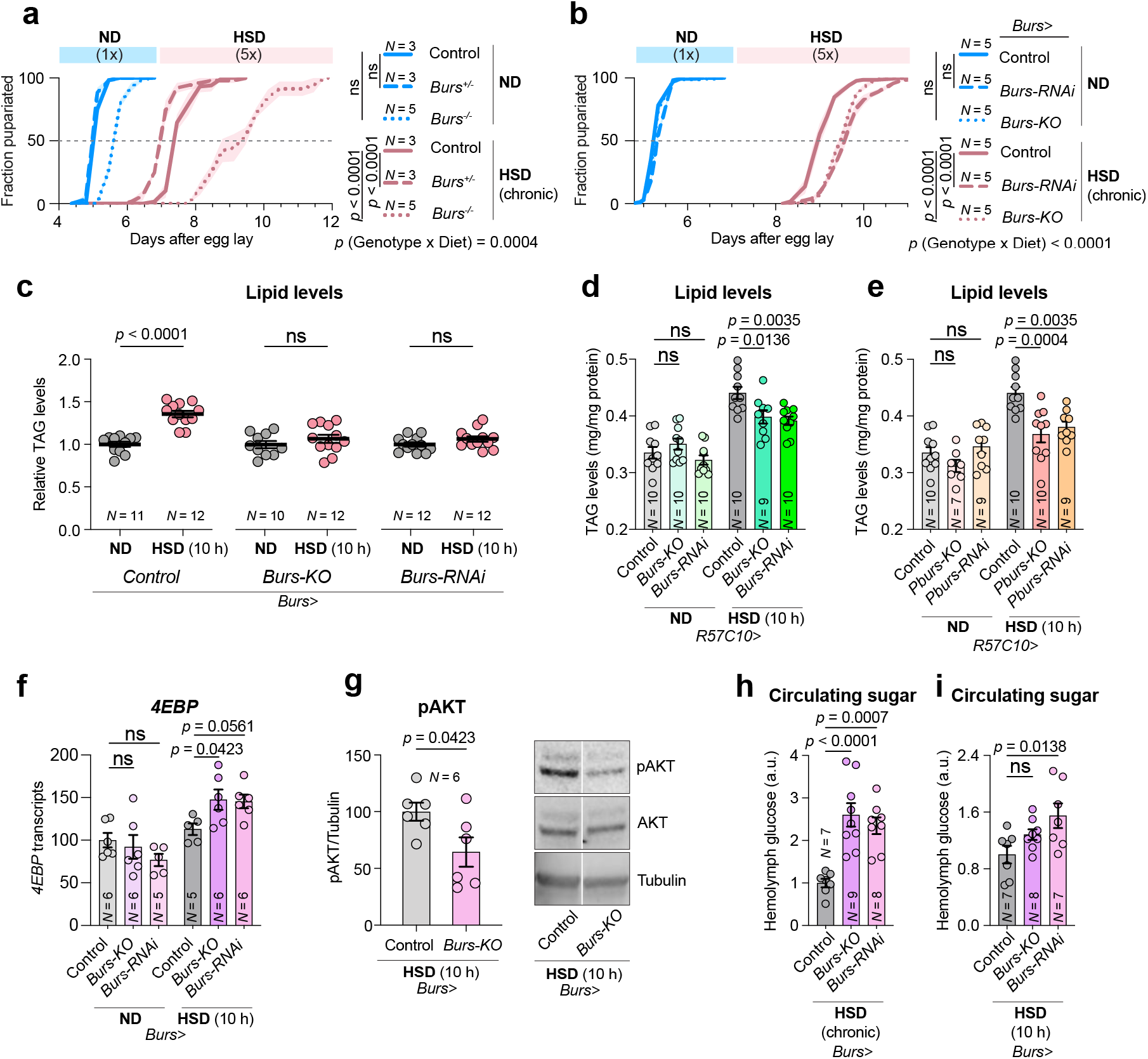
Both Bursicon subunits – Burs and PBurs – are required for metabolic and developmental adaptation to a high-sugar diet. (A) Pupation timing on normal and high-sugar diets for controls and *Burs^Z4410^* null-mutant heterozygotes and homozygotes. (B) Pupation timing on normal and high-sugar diets for animals expressing knockdown or knockout of *Burs* in the *Burs*-expressing cells (with *Burs-GAL4*). See also Extended Figure 2A. (C) Stored lipid levels in animals raised for 90 hours on a normal diet and then fed a 10-hour pulse of normal or high-sugar diet, comparing controls and animals expressing *Burs* knockdown or knockout in the *Burs*-expressing cells. (D Stored lipid levels in animals raised for 90 hours on a normal diet and then fed a 10-hour pulse of normal or high-sugar diet, comparing controls and animals expressing *Burs* knockdown or knockout in all neurons. (E) Stored lipid levels in animals raised for 90 hours on a normal diet and then fed a 10-hour pulse of normal or high-sugar diet, comparing controls and animals expressing *Pburs* knockdown or knockout in all neurons. (F) Whole-body *4EBP* transcript levels measured by qPCR in controls and animals with *Burs* loss in the *Burs*-expressing cells, raised on normal diet for 90 hours and then fed with a 10-hour pulse of normal or high-sugar diet. (G) Whole-body pAkt levels, normalized to alpha-Tubulin, measured by Western blot, in controls and *Burs*-cell *Burs*-deletion animals fed high-sugar diet for ten hours after 90 hours on normal medium. (H) Hemolymph glucose levels of control and *Burs*-cell *Burs*-loss animals raised on high-sugar diet. (I) Hemolymph glucose levels of control and *Burs*-cell *Burs*-loss animals raised on a normal diet and exposed to a 10-hour pulse of high-sugar diet. Statistics: Error bars represent mean and SEM. ns, not significant (*p*>0.05). A, one-way Kruskal-Dunn ANOVA for multiple comparison between 50%-pupariation times and two-way ANOVA for interaction. B, one-way Tukey ANOVA for multiple comparison between 50%-pupariation times and two-way ANOVA for interaction. C, Mann-Whitney nonparametric U test between diets. D, E, F, one-way ANOVAs with Dunnett’s correction. G, two-sided unpaired parametric t-test. H, one-way ANOVA with Dunnett’s correction. I, one-way ANOVA with Dunnett’s correction.

The capacity to adapt to conditions of high sugar intake directly depends on the ability to store excess energy by converting it into fat that is deposited in adipose tissues. Animals raised on HSD, therefore, exhibit increased body fat stored as triacylglycerides (TAGs)^8,9^. We therefore investigated whether Bursicon signaling mediates metabolic adaptation to nutrient excess by assessing animals’ fat storage. We observed a significant increase in whole-body TAG levels in animals fed a high-sugar diet for 10 hours, an effect that was abolished by *Burs* loss (Fig. 2c). This effect was also attenuated in animals expressing neuronal RNAi or CRISPR targeting *Burs* or *Pburs*, whereas on ND these animals’ TAG levels were unchanged from those of controls (Fig. 2d,e). This indicates that Bursicon signaling is required for excess glucose to be converted to fat for storage on high-sugar diets.

Insulin resistance impairs the cellular uptake of glucose and its conversion to glycogen and fat for storage. We therefore assessed the possibility that the lipid-storage defect observed in animals with reduced neuronal Burs expression might arise from impaired insulin signaling. Inhibition of Bursicon signaling through neuronal *Burs* knockdown or knockout led to increased whole-body expression of *4EBP*, a target gene of FOXO whose expression is repressed by insulin signaling^34^, on HSD but not on ND (Fig 2f), suggesting decreased peripheral insulin-signaling. Consistent with this observation, neuronal loss of *Burs* was also associated with decreased phosphorylation of the insulin-signaling component AKT in whole-animal samples (Fig. 2g), collectively suggesting that loss of *Burs* expression impairs insulin signaling on high-sugar diet. Impairments in insulin signaling reduce tissue glucose uptake from circulation, leading to hyperglycemia^8^. Consistent with reduced insulin signaling, loss of *Burs* led to elevated glycemic levels after chronic or short-term (10-hour) exposure to a high-sugar diet (Fig. 2h,i). Together, our results indicate that neuronal Bursicon signaling regulates insulin signaling or sensitivity to adapt systemic metabolism and maintain glycemic homeostasis in response to a high-sugar diet.

### Burs^+^ neurons respond to dietary sugar and control insulin production

Insulin is the key regulatory hormone that drives the reduction of blood glucose levels after sugar intake. Secretion of insulin from the insulin-producing cells (IPCs) – the main source of circulating insulin in *Drosophila* and analogous with the pancreatic β-cells – is governed by many nutritional cues from peripheral organs^34,35^. To test whether Bursicon signaling regulates insulin secretion on HSD, we assessed the effect of *Burs* loss on the levels of insulin-like peptides 2, 3 and 5 (Ilp2, −3, and −5), the three main insulins produced by the IPCs. Consistent with the increased *4EBP* expression and lower phosphorylated-AKT levels in animals with reduced Burs signaling on HSD, which suggest reduced systemic insulin signaling on HSD (Fig. 2f,g), we observed a strong reduction in *Ilp2* and *Ilp3* transcript levels, albeit without any change in *Ilp5* expression, in the central nervous system (CNS) of animals with loss of *Burs* function, indicating reduced insulin production (Fig. 3a). Consistent with this lower expression, the levels of Ilp2 and Ilp3 peptides, but not of Ilp5, were reduced in the IPCs (Fig. 3b and Extended data Fig. 3a) in animals with loss of *Burs*. Taken together, these findings suggest that, in response to consumption of dietary sugar, Bursicon signaling acts on the IPCs to promote the production of Ilp2 and Ilp3 to increase insulin signaling and enhance tissue uptake of circulating glucose.

**Figure 3.**
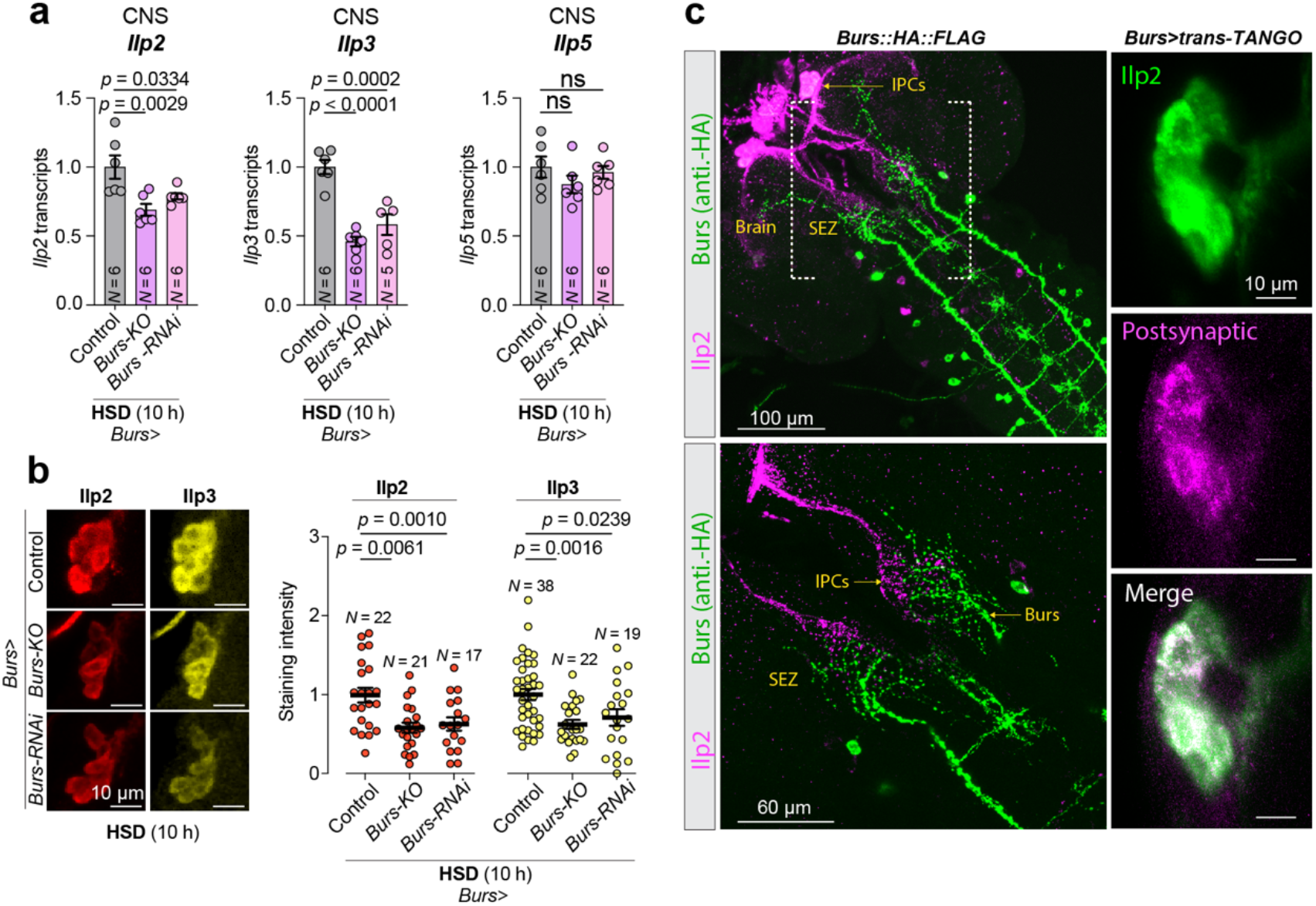
Burs regulates the IPCs, likely through direct neuronal communication. (A) Transcript levels of *Ilp2*, −*3*, and −*5* in dissected CNS samples of controls and animals with *Burs*-cell *Burs* loss, exposed to high-sugar diet for 10 hours after 90 hours’ feeding on normal diet. (B) Left: Representative images showing levels of Ilp2 and Ilp3 retained within the IPCs in animals with Burs^+^ cell *Burs* loss. Right: Quantified levels of these peptides in IPCs from individual animals. (C) Left: Larval CNS stained for Ilp2 (purple) and HA::Burs::FLAG (anti-HA, green). Right: *Burs*-cell-originating trans-Tango signal (purple) in the IPCs, marked with anti-Ilp2 (green). Scale bars: Left: top, 100 microns; bottom, 60 microns. Right: 10 microns. Statistics: Error bars represent mean and SEM. ns, not significant (*p*>0.05). A, one-way ANOVAs with Dunnett’s correction. B, one-way ANOVA with Dunnett’s correction.

We noticed that Burs^+^ neurons send projections to the sub-esophageal zone (SEZ), a region in which the IPCs also arborize. To examine the proximity of arbors of Burs+ neurons and the IPCs, we co-stained brains for both neuronal populations and found that neurites of the Burs^+^ neurons project in close proximity to the arbors of the IPCs (Fig. 3c), suggesting that the IPCs might receive input from Burs^+^ neurons. We therefore sought to test for synaptic connectivity between the two neuronal populations using the anterograde trans-synaptic tracing technique trans-Tango, in which a tethered ligand expressed in pre-synaptic cells activates its cognate receptor in postsynaptic target neurons, leading to *tdTomato* expression in these cells^36^. We observed a trans-Tango-dependent *tdTomato* signal in the IPCs when the presynaptic ligand was expressed in Burs^+^ neurons (Fig. 3c), indicating that the IPCs are directly postsynaptic to the Burs^+^ neurons.

To assess whether Bursicon itself acts directly on the IPCs, we examined whether the larval IPCs express the Burs:Pburs receptor Rickets (Rk) using a T2A::GAL4 knock-in into the endogenous *rk* locus to drive expression of *UAS-mCD8::GFP*. We observed GFP reporter expression in the IPCs (Fig. 4a), indicating that *rk* is expressed in these cells. This is further supported by single-nucleus transcriptomics^37^, together suggesting that the IPCs are receptive for Bursicon signaling. Next, we investigated whether loss of Bursicon signaling in the IPCs might be responsible for the HSD-induced developmental delay observed in Burs-/Pburs-loss animals. We found that knockdown or knockout of *rk* in the IPCs resulted in an HSD-induced developmental delay (Fig. 4b) along with reduced *Ilp2* and *Ilp3* transcript levels, with unchanged expression of *Ilp5* (Fig. 4c), phenocopying *Burs* loss. The reduced *Ilp* transcript levels were also reflected by reduced levels of Ilp3 peptides in the IPCs (Fig. 4d). Consistent with this, loss of Rk activity in the IPCs led to reduced levels of circulating levels of Ilp2 peptide in the hemolymph (Fig. 4e). The activation of Rk by the Burs:Pburs heterodimer leads to increased production of cAMP^31^, a second messenger that in mammalian beta cells promotes glucose-stimulated insulin secretion^38^. Since many effects of cAMP are mediated by cAMP-dependent protein kinase (PKA), we investigated the impact of blocking PKA signaling in the IPCs. Inhibition of PKA through expression of a dominant-negative (cAMP-insensitive) form of the regulatory subunit R1 in the IPCs led to HSD-dependent delay, similar to that observed with loss of neuronal *Burs/Pburs* or IPC-specific loss of *rk* (Fig. 4f). This indicates that the insulinotropic effects of signaling from Burs neurons to the IPCs, through Rk, are at least in part mediated by cAMP/PKA signaling. In contrast to the effects of neuronal *Burs* or *Pburs* loss, IPC-specific knockdown or knockout of *rk* did not lead to changes in TAG levels on HSD condition (Fig. 4g), indicating that neuronal Burs signaling regulates fat storage via tissues other than the IPCs.

**Figure 4.**
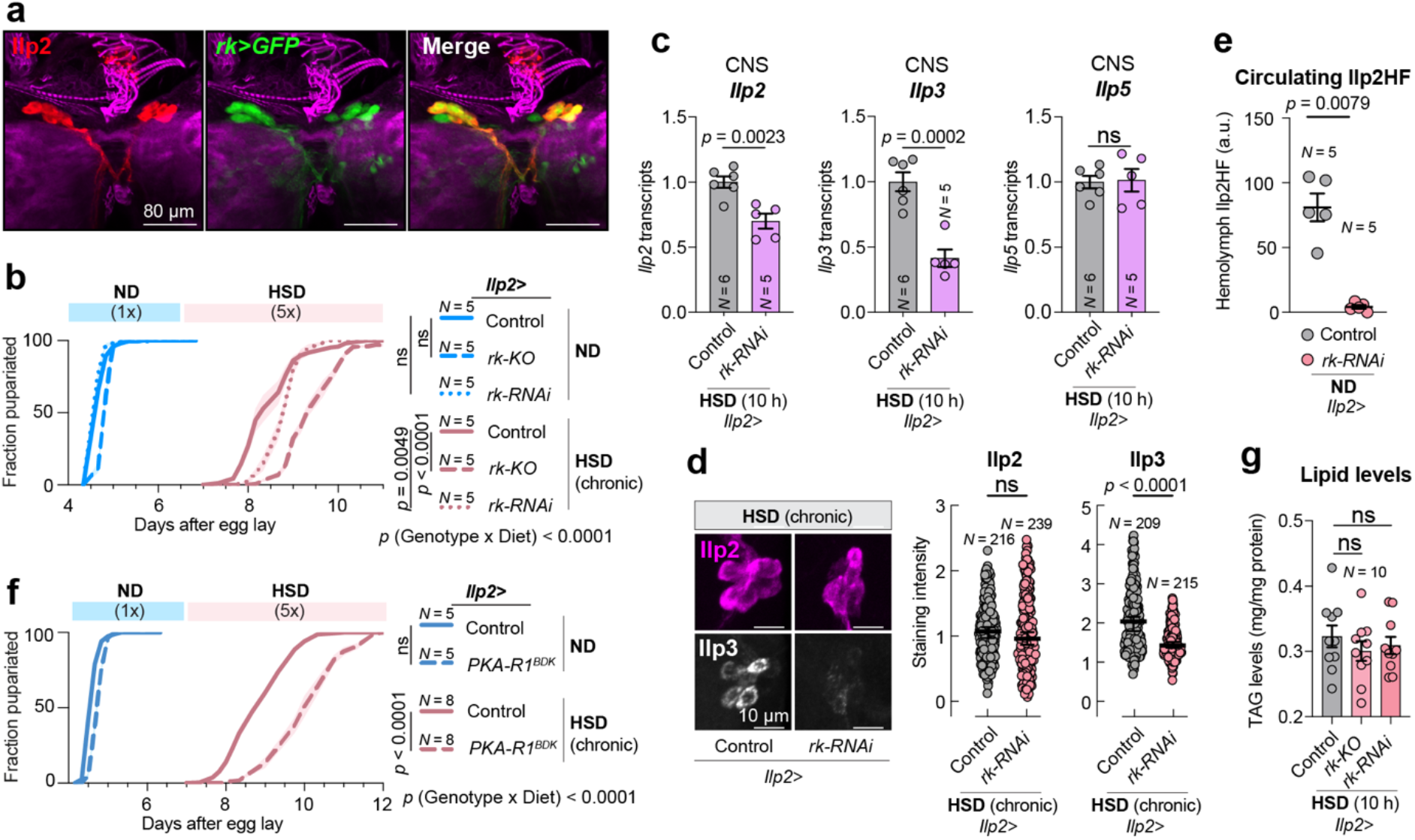
Rickets-mediated signaling regulates the IPCs and phenocopies Burs manipulations. (A) Larval brain stained for Ilp2 (red), *rickets-GAL4>GFP* (green), and filamentous actin (purple). Scale bars, 80 microns. (B) Pupation timing of controls and animals with IPC-specific *rickets* knockdown or deletion, on normal and high-sugar diet. (C) Transcript levels of *Ilp2*, −*3*, and −*5* in dissected CNS samples of animals with *IPC*-specific *rickets* knockdown raised normal diet for 90 hours and fed 5x-sugar for 10 hours. (D) Left: Representative images showing levels of Ilp2 and Ilp3 retained within the IPCs in controls and animals with IPC-specific *rickets* knockdown, raised on high-sugar diet. Right, quantification of these peptide levels in multiple samples. Scale bars at left, 10 microns. (E) ELISA against circulating tagged Ilp2 in controls and animals expressing RNAi against *rickets* in the IPCs, raised on normal diet. (F) Pupation timing of controls and animals with IPC-specific expression of dominant-negative PKA regulatory subunit (PKA-R1^BDK^), raised on normal or high-sugar diet. (G). Whole-larval lipid-storage levels of controls and animals lacking *rickets* in the IPCs, raised on normal diet for 90 hours and transferred to high sugar for 10 hours. Statistics: Error bars represent mean and SEM. ns, not significant (*p*>0.05). B: one-way ANOVA with Tukey’s correction for multiple comparison between 50%-pupariation times and two-way ANOVA for interaction. C, two-sided unpaired t-tests. D, Mann-Whitney nonparametric U tests. E, Mann-Whitney nonparametric U test. F, two-sided unpaired t-test between 50% times and two-way ANOVA for interaction. G, one-way ANOVA with Tukey’s correction for multiple comparison.

### Systemic Bursicon signaling to fat cells protects against insulin resistance to maintain glucose homeostasis on high sugar diets

In addition to its signaling within the CNS, Bursicon (Burs:Pburs) hormone is released into the circulation from neurohemal release sites^39^, suggesting a possible humoral role of Bursicon in metabolic adaptations to high sugar. To identify the target tissues mediating any effect of humoral Bursicon signaling on fat storage, we assessed the tissue expression of *rk* and found strong expression in the fat body, an adipose-like organ that stores excess energy as TAG and glycogen. This receptor expression suggested that direct action of Bursicon on the adipose cells might cause the observed lipid-storage effects of *Burs/Pburs* loss. Therefore, we investigated whether Rk in the fat body is involved in Bursicon-mediated regulation of sugar tolerance. Knockdown or knockout of *rk* in the larval fat body indeed led to an HSD-induced delay (Fig. 5a), suggesting that loss of Bursicon signaling in the fat body impairs glucose tolerance. We then assessed whether this sugar intolerance was associated with an inability to store excess energy as fat. Loss of *rk* in the fat body had no effect on lipid levels on a normal diet, but this manipulation significantly attenuated the increase in TAG associated with high-sugar feeding (Fig. 5b). In mammalian and *Drosophila* adipose tissue, stored fat is deposited within specialized lipid droplet organelles. We therefore investigated the effect of fat-body Bursicon/Rickets signaling on lipid storage droplets in response to high-sugar feeding. Consistent with the elevated TAG levels observed in these conditions, we found that high-sugar feeding increased the area of fat-body cells that was occupied by lipid droplets in control animals (Fig. 5c,d), indicating increased fat storage. In animals with *rk* knockdown, however, the lipid-droplet area remained unchanged in response to high-sugar feeding, indicating that loss of Bursicon signaling in the adipose tissue is associated with an inability of the tissue to process the excess energy and store it as fat.

**Figure 5.**
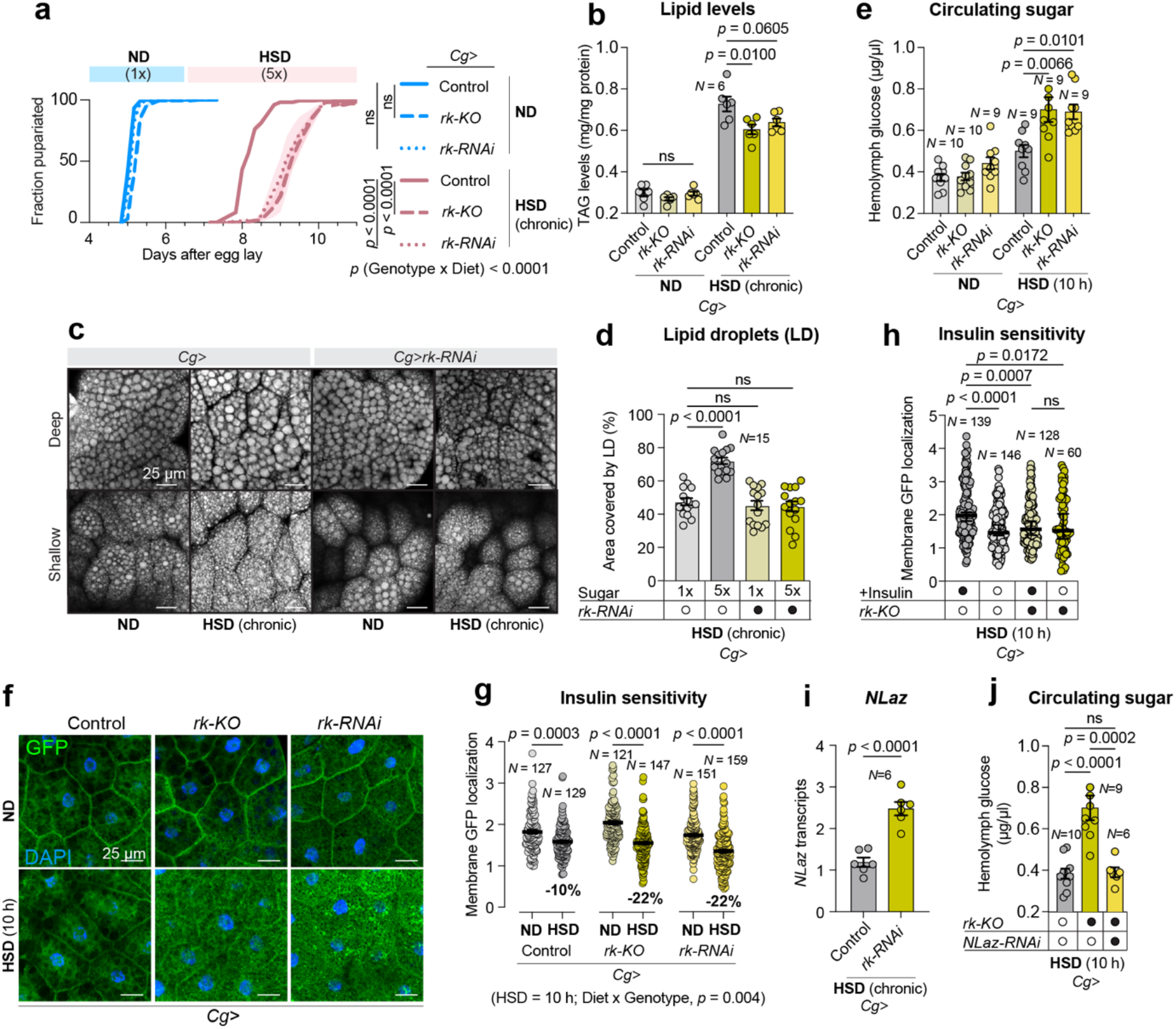
Rickets in the fat body systemically regulates development and metabolism. (A) Pupariation timing of controls and animals expressing fat-body-specific *rickets* knockdown or deletion, raised on normal or high-sugar diet. (B) Lipid-storage levels of controls and animals with fat-body *rickets* loss, raised on normal and high-sugar diets. (C) Images of lipid droplets, stained with Nile Red, in fat-body tissues of controls and animals lacking *rickets* in the fat body, raised on normal or high-sugar diet. Top, a plane at the middle of the tissue; bottom, a more superficial, glancing plane. Scale bars, 25 microns. (D) Quantification of tissue area containing lipid in these animals. (E) Circulating glucose levels of controls and animals lacking *rickets* in the fat body, raised for 90 hours on normal diet and transferred to either normal or high-sugar diet for 10 hours. (F) Representative images of the tGPH *in-vivo* insulin-signaling indicator in controls and animals lacking fat-body *rickets* expression. Animals were raised for 90 hours on normal diet and transferred to fresh normal medium (top) or 5x-sugar medium (bottom) for 10 hours. Scale bars, 25 microns. (G) The ratio of at-membrane to near-membrane GFP signal, a quantification of tGPH localization, from multiple samples of these animals. (H) The ratio of at-membrane to near-membrane GFP signal in fat-body explants from controls and animals lacking fat-body *rickets,* raised for 90 hours on normal food and then transferred to high-sugar food for 10 hours, and either stimulated with human insulin or not. (I) The level of *NLaz* transcripts measured by qPCR in whole control larvae or animals expressing fat-body *rickets* loss, raised on high-sugar diet. (J) Hemolymph glucose levels of controls, animals expressing *rickets* deletion in the fat body, and animals expressing both *rickets* deletion and *NLaz* knockdown in the fat body, all raised for 90 hours on normal diet and then transferred to high-sugar medium for ten hours. Statistics: Error bars represent mean and SEM. ns, not significant (*p*>0.05). A, one-way Kruskal-Wallis nonparametric ANOVA with Dunn’s correction between 50%-pupariation times for multiple comparisons and two-way ANOVA for interaction. B, one-way ANOVA with Dunnett’s correction. D, one-way ANOVA with Dunnett’s correction. E, one-way ANOVA with Dunnett’s correction. G, two-way ANOVA and one-way ANOVA with Tukey’s correction. H, one-way Kruskal-Wallis nonparametric ANOVA. I, two-sided unpaired t-test. J, one-way ANOVA with Tukey’s correction.

These results suggest that Bursicon signaling through its cognate LGR receptor Rickets is important for the absorption of excess circulating glucose by the fat body or for the conversion of intracellular carbohydrates into triglycerides for storage. We therefore examined whether fat-body Bursicon/Rickets signaling might affect circulating sugar concentrations and found that *rk* loss in the fat body led to HSD-induced hyperglycemia (Fig. 5e), consistent with impaired glucose uptake into the fat tissue. This hyperglycemic effect is often caused by insulin resistance, which reduces insulin-stimulated glucose uptake into tissues. To examine whether loss of Bursicon signaling in adipocytes impairs insulin signaling, we measured the responsiveness of fat-body tissues to insulin using tGPH, a fluorescent indicator that comprises a GFP moiety fused to a PI(3)P-binding Pleckstrin-homology domain and thus reflects insulin/PI3K activity by increased membrane association^40^. Fat-body cells with *rk* loss showed a stronger reduction in cell-membrane localization of GFP in response to high-sugar feeding compared to controls, indicating that inhibition of Bursicon signaling in adipose tissue reduces insulin/PI3K activity on high-sugar diet (Fig. 5f,g). These results indicate that loss of Bursicon signaling in fat cells leads to reduced basal insulin signaling – either central production or peripheral sensitivity – in HSD conditions. To identify whether insulin sensitivity was reduced in adipose tissue with loss of Bursicon signaling, we measured membrane GFP localization in adipocytes in response to *ex-vivo* stimulation with human insulin. Fat tissue from control animals fed HSD responded to exogenous insulin with an increase in insulin/PI3K activity and thus membrane GFP localization, whereas fat bodies with *rk* knockdown showed no response to this insulin stimulation (Fig. 5h). Taken together, these observations suggest that fat-body loss of Bursicon signaling results in systemic glucose intolerance and lack of glycemic control, coupled with adipocyte insulin resistance under high-sugar-diet conditions.

Insulin resistance is a main component of the pathogenesis of diabetes, and the mechanisms involved include cellular stress and inflammatory responses. The upregulation of expression of the lipocalin Neural Lazarillo (NLaz, mammalian ApoD) through the Jun-N-terminal kinase (JNK) pathway is proposed to be a part of the cellular stress-response pathway that contributes to the development of insulin resistance in HSD – indeed, NLaz is necessary and sufficient for this effect^9^. We therefore assessed whether Bursicon signaling affects *NLaz* expression in the fat tissue. Knockdown of *rk* in the fat body led to *NLaz* upregulation in whole-animal assays (Fig. 5i), consistent with insulin resistance. We then tested the ability of *NLaz* deficiency to rescue the defect in glycemic control induced by *rk* loss in HSD conditions and found that reducing *Nlaz* expression in the fat alone completely alleviated the hyperglycemia caused by *rk* knockdown in that tissue under high-sugar conditions (Fig. 5j). These data place NLaz downstream of Burs/Rickets, indicating that Burs/Rickets activity normally inhibits NLaz expression on HSD, and show that this inhibition is indeed critical for maintaining glycemic control.

### Inhibition of NF-κΒ protects adipocytes against insulin resistance during high-sugar feeding

In addition to its metabolic functions, the fat body is also the main immune-responsive organ in the fly. Immune pathways are highly integrated with metabolic programs to ensure sufficient energy to support immune responses during infection. The fat body expresses the conserved immune-responsive transcription factor Nuclear Factor κB (NF-κB), named Relish (Rel) in the fly^41^. Relish has been placed downstream of Burs:Burs and PBurs:PBurs homodimers, but not Rickets, in adult fat-tissue immune responses^42^. We therefore explored the possibility that interaction between Relish signaling and metabolic-control pathways might underlie the observed sugar-induced phenotypes. If Relish mediates effects of Burs/Rk signaling in the fat body, one would expect that loss of this factor would phenocopy neuronal *Burs/Pburs* loss and fat-body *rk* loss and exacerbate HSD-induced delay. Surprisingly, we found the opposite effect: fat-body specific *relish* knockdown partially rescued the HSD-induced developmental delay (Fig. 6a). This indicates that inhibition of *relish* alleviates high-sugar-induced metabolic defects in adipose tissues and suggests that Relish does not mediate the effects of Burs/Pburs/Rk on sugar tolerance. This is consistent with findings showing that the Relish-mediated immune responses to Bursicon signaling are driven by Burs and Pburs homodimers through a Rk-independent mechanism, likely via interactions with an unknown receptor^42^. Our findings indicate that Rk-mediated Bursicon-heterodimer signaling and NF-κB/Relish signaling in the adipose tissue have opposing effects on sugar tolerance and potentially implicate the activation of the NF-κB pathway in the pathology of diabetes in response to overnutrition.

**Figure 6.**
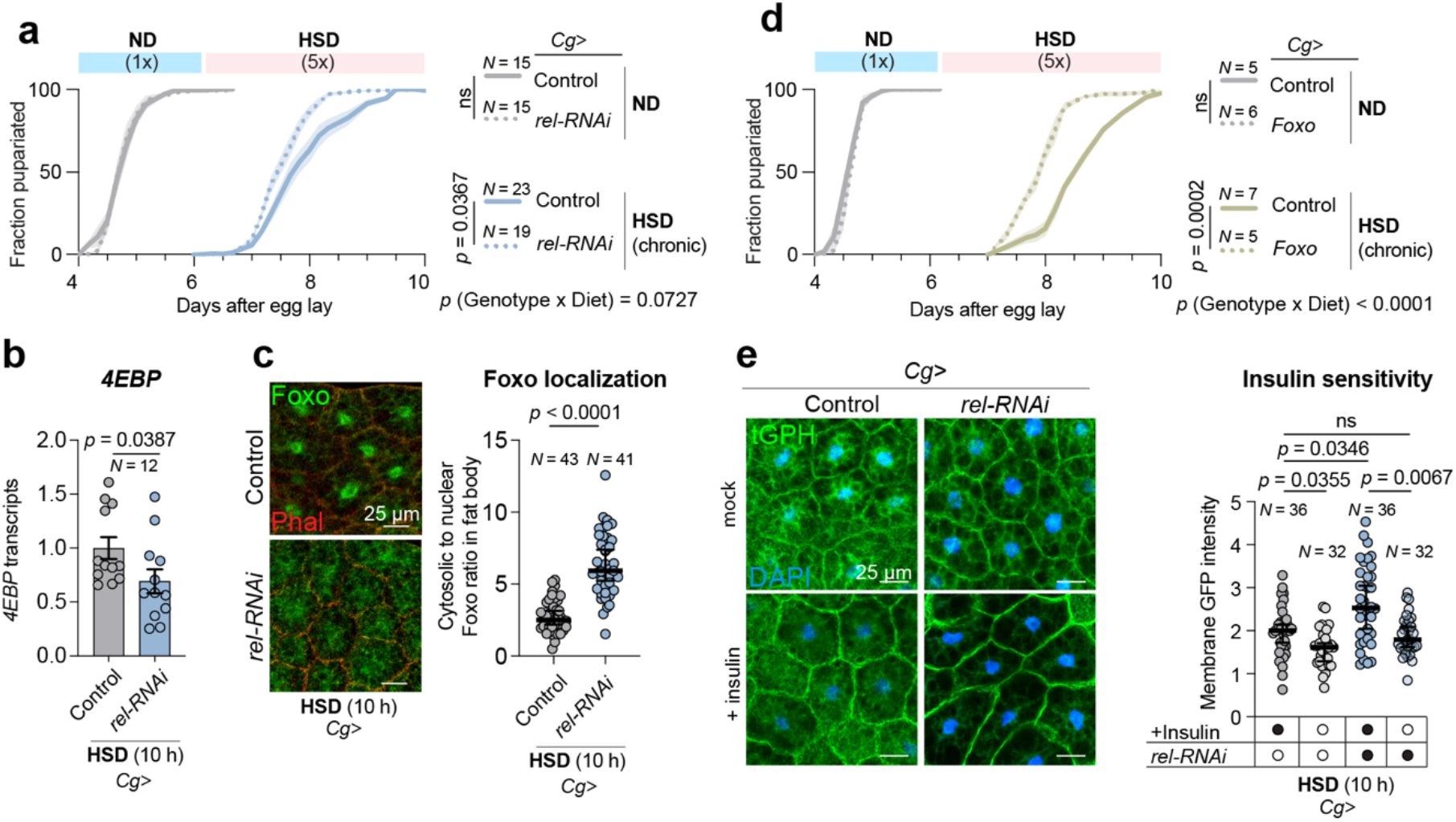
Relish (NF-κB) loss in the fat body counteracts high-sugar-diet-induced effects. (A) Pupariation timing of controls and animals expressing fat-body-specific *relish* knockdown, raised on normal or high-sugar diet. (B) Transcript levels of *4EBP* measured by qPCR in carcass samples of controls and fat-body *relish* knockdowns raised on normal diet for 90 hours, followed by 10 hours of high-sugar medium. (C) Left: representative images of Foxo localization (green; filamentous actin costained in red) in fat-body tissue from controls and animals expressing fat-body *relish* knockdown, after 10-hour high-sugar feeding. Right: a quantification of Foxo distribution, the ratio of cytoplasmic staining to nuclear staining. Scale bars, 25 microns. (D) Pupariation timing of controls and animals overexpressing *Foxo* in the fat body, raised on normal and high-sugar diet. (E) Left: representative images showing tGPH insulin-indicator activity in fat-body tissues of controls and animals expressing fat-body *relish* knockdown, fed high-sugar diet for 10 hours; explants were stimulated with control solution or with human insulin. Scale bars, 25 microns. Right: Center-to-surround ratio of tGPH membrane signal for these animals. Statistics: Error bars represent mean and SEM. ns, not significant (*p*>0.05). A, two-sided unpaired t-tests between 50%-pupariation times for multiple comparisons and two-way ANOVA for interaction. B, Mann-Whitney nonparametric U test. C, Mann-Whitney nonparametric U test. D, two-sided unpaired t-tests between 50%-pupariation times for pair-wise comparisons and two-way ANOVA for interaction. E, Kruskal-Wallis nonparametric ANOVA.

We therefore asked whether NF-κB/Relish inhibition in the adipose tissue protects against diet-induced insulin resistance. In general, increased insulin signaling induces the phosphorylation of the transcription factor FOXO, causing its exclusion from the nucleus, which leads to downregulation of the FOXO transcriptional target *4EBP*^34^. We observed that, in animals fed a high-sugar diet, knockdown of *relish* in the fat body led to reduced nuclear localization of FOXO (Fig. 6c) and an associated reduction in *4EBP* expression (Fig. 6b), indicating increased insulin signaling in the fat tissue. Thus, reduction of NF-κB/Relish activity inhibits FOXO and leads to a partial reversal of the sugar-induced diabetic phenotype, suggesting that HSD-induced metabolic phenotypes and insulin resistance are associated with increased activity of FOXO. Consistent with this, overexpression of *FOXO* exacerbated the HSD-induced developmental delay, indicating that FOXO is a mediator of sugar-induced insulin resistance (Fig. 6d).

Our findings indicate that inhibition of NF-κB/Relish signaling reverses sugar-induced insulin-resistance phenotypes. To test directly whether this pathway affects insulin sensitivity, we stimulated adipocytes from HSD-fed animals with human insulin *ex vivo*. In agreement with increased insulin sensitivity, the effect of insulin stimulation on insulin-PI3K reporter activity (tGPH membrane-associated GFP) was larger in adipose tissue from *relish*-knockdown animals than in tissue from control animals, suggesting increased insulin sensitivity (Fig. 6e). This suggests that inhibition of NF-κB/Relish ameliorates diet-induced insulin resistance, opening the interesting possibility that this immune pathway can be targeted for the reversal of diabetes.

### Bursicon mediates a muscle-neuronal-adipose relay that controls insulin signaling and adipocyte insulin sensitivity

Since neuronal Bursicon signaling is required for sugar tolerance, we investigated whether Burs^+^ neurons respond to dietary sugar. Burs is expressed in pairs of neurons in the subesophageal zone (SEZ) and the thoracic and abdominal segments in the larval CNS, whereas the expression of Pburs is confined to a subset of Burs^+^ neurons in the abdominal segments^31^. Consequently, the production of the active heterodimeric Bursicon hormone within the CNS is restricted to those abdominal neurons that express both the Burs and Pburs subunits. We used the CaLexA system^43^ to explore whether high-sugar feeding regulates calcium activity in the bilateral neurons in the thoracic (T3) and abdominal segments (A1-4) that co-express Burs and Pburs^31^. Strong calcium-induced GFP signal was observed solely in Burs^+^ neurons that produce the Burs:Pburs heterodimer, indicating that these are the active Burs^+^ neurons (Fig. 7a). We then quantified calcium activity of these neurons in response to dietary sugar. Chronic high-sugar feeding increased calcium signaling but did not affect intracellular Burs peptide levels in these cells, demonstrating that the subset of neurons producing the Burs/Pburs heterodimer are responsive to dietary sugar (Fig. 7b-d). Conversely, Burs peptide levels were reduced in the SEZ and thoracic pairs of neurons that do not express Pburs (Fig.7e), indicating that neurons that might produce the possible Burs homodimer are affected differently by dietary sugar feeding.

**Figure 7.**
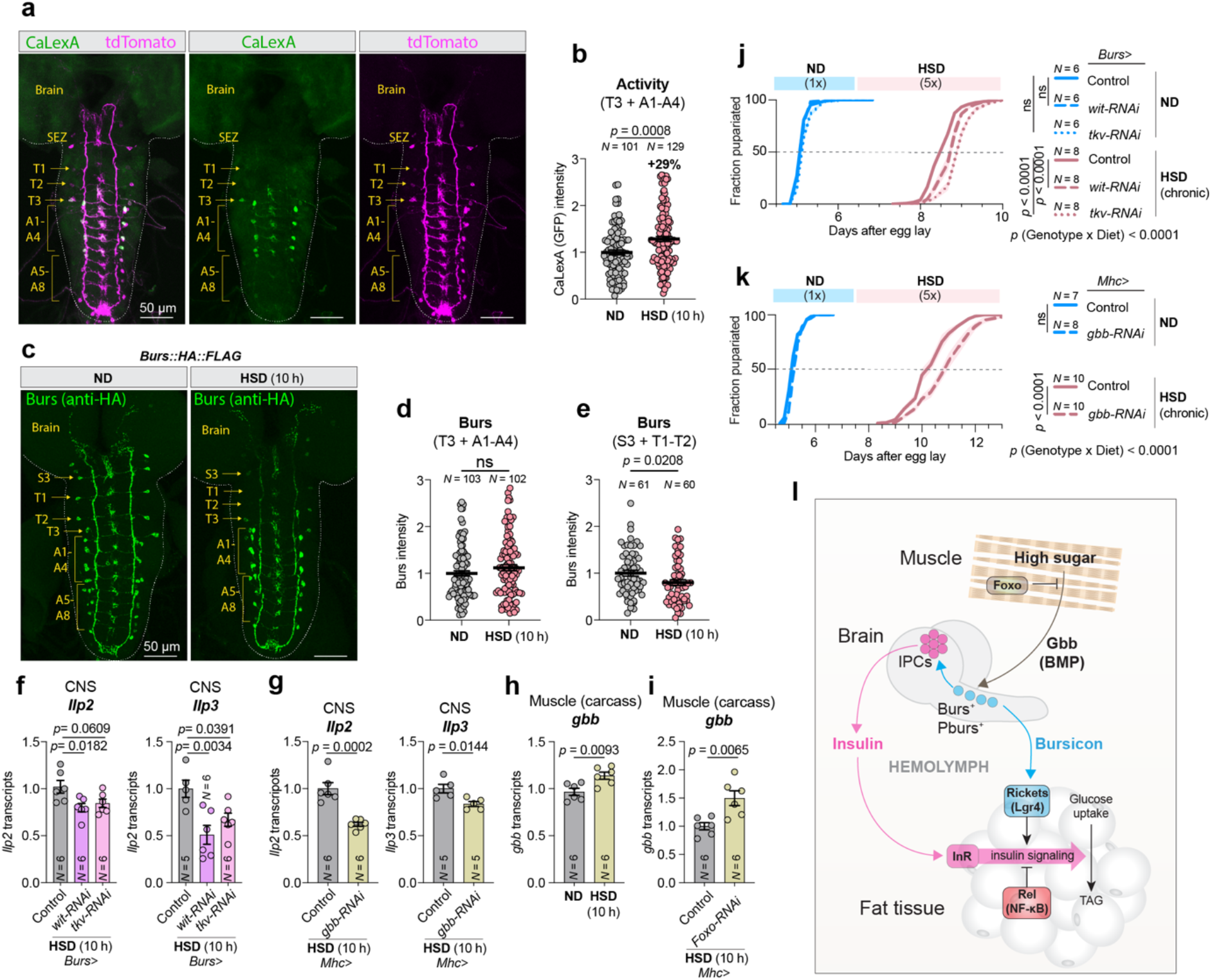
High sugar activates a subset of Burs-expressing cells – Pburs-coexpressing cells that project to the larval musculature – via retrograde BMP signaling from the musculature. (A) Representative images of larval CNS from an animal expressing CaLexA (green) and tdTomato (purple) in the *Burs*-expressing cells. Segments within the CNS are labeled, in animals raised on 1x sugar for 90 hours and transferred to 5x-sugar food for 10 hours. Scale bars, 50 microns. (B) Quantification of CaLexA signal (GFP) in *Burs*-expressing cells of segments T3 and A1-A4, normalized to tdTomato intensity, in animals raised on 1x sugar for 90 hours and transferred to either fresh 1x medium or 5x-sugar food for 10 hours. (C) Representative images of HA::Burs::FLAG peptide levels (anti-HA, green) in control animals fed normal or high-sugar diet for 10 hours. Segments within the CNS are labeled. Scale bars, 50 microns. (D) Quantification of Burs staining intensity in segments T3 and A1-A4 of brains from animals fed normal or high-sugar diet for 10 hours. (E) Quantification of Burs staining intensity in segments S3, T1, and T2 of the same brains. (F) Transcript levels of *Ilp2* and *Ilp3* measured by qPCR in brain samples from controls and animals expressing RNAi against *wit* or *tkv* in the *Burs*-expressing cells, fed high-sugar medium for 10 hours. (G) Transcript levels of *Ilp2* and *Ilp3* measured by qPCR in brain samples from controls and animals expressing RNAi against *gbb* in the musculature, fed high-sugar medium for 10 hours. (H) Transcript levels of *gbb* measured by qPCR in carcass samples from control animals fed for 10 hours with normal or high-sugar medium. (I) Transcript levels of *gbb* measured by qPCR in carcass samples from controls and animals expressing muscle-specific RNAi against *Foxo*, fed high-sugar diet for 10 hours. (J) Pupariation timing of controls and animals lacking *wit* or *tkv* in the *Burs*-expressing cells, raised on normal or high-sugar diet. (K) Pupariation timing of controls and animals lacking *gbb* expression in the musculature, raised on normal or high-sugar diet. (L) A model consistent with the findings of this work. Statistics: Error bars represent mean and SEM. ns, not significant (*p*>0.05). B, D, E, Mann-Whitney nonparametric U tests. F, one-way ANOVA with Dunnett’s correction. G, H, I, two-sided unpaired t-tests. J, K, one-way ANOVAs with Tukey’s correction between 50%-pupariation times for multiple comparisons and two-way ANOVA for interaction.

The BMP ligand Glass bottom boat (Gbb) has previously been shown to activate neuronal expression of Pburs^44^. Muscle-derived Gbb activates the TGF-β receptors Wishful thinking (Wit) and Thickveins (Tkv) in Burs^+^ neurons, providing a retrograde signal from the muscle that upregulates neuronal Pburs. Notably, Tkv and other components of the TGF-β pathway were identified as a hits in our screen for genes involved in sugar tolerance (Fig. 1e,d). We therefore examined whether muscle-derived Gbb regulates neuronal Bursicon signaling under high-sugar*-*feeding conditions. We found that knockdown of *gbb* in the muscles or of *tkv* in the Burs^+^ neurons reduced the expression of *Pburs* in the CNS (Extended Fig. 4a), confirming previous observations that muscle-derived Gbb acts in a retrograde manner to regulate Bursicon signaling in the CNS. We then examined the role of Gbb in the muscle as a potential peripheral regulator of central insulin signaling in the nervous system. Knockdown of *gbb* in the muscles or of its receptor *tkv* in the Burs^+^ neurons resulted in reduced expression of *Ilp2* and *Ilp3*, similar to the effects observed in animals with neuronal loss of *Burs* or *Pburs* and in those with loss of *rk* in the IPCs themselves (Fig. 3a, 4c, 7f,g). This suggests that muscles are the source of Gbb that regulates insulin through Bursicon in response to high-sugar feeding. We next examined whether Gbb production in the musculature is regulated by high-sugar feeding by measuring *gbb* transcript levels in dissected body-wall muscles included in the carcass. *Gbb* expression was mildly but significantly upregulated in muscles in response to a chronic HSD (Fig. 7h).

Dietary sugar can inhibit FOXO activity in muscles through insulin-mediated signaling pathways. We therefore wondered whether reducing FOXO action in the muscle would promote expression of *gbb*. To test this hypothesis, we knocked *FOXO* down in the musculature and assessed the expression of *gbb*. We found an upregulation of muscle *gbb* expression when *FOXO* was knocked down during high-sugar feeding (Fig. 7i), indicating that FOXO inhibits Gbb expression in the musculature. This suggests that FOXO in the muscle is a dietary-sugar-regulated effector that governs Gbb/BMP signaling. Therefore, our results indicate that in response to high-sugar feeding, FOXO-regulated muscle BMP/Gbb signaling remotely regulates insulin production in the IPCs through a neuronal relay mediated by Bursicon signaling. We then investigated whether this muscle-neuronal relay is important for sugar tolerance by examining the effect on HSD-induced phenotypes of silencing *gbb* in the muscles or its receptors in Bursicon neurons. We found that either muscle-specific loss of *gbb* or knockdown of its receptors *wit* or *tkv* in the Burs^+^ neurons resulted in a HSD-induced delay (Fig. 7j,k), mimicking the phenotype observed with loss of neuronal *Burs*/*Pburs* and IPC- or fat-body-specific loss of *rk* (Fig. 2a,b, 4b, and Extended data Fig. 2a-c). Collectively, our results indicate that high-sugar feeding increases muscle-derived Gbb signaling onto Burs^+^/Pburs^+^ neurons of the CNS, leading to increased Bursicon signaling directly to the IPCs, which stimulates insulin production, and to the fat body, which enhances its insulin sensitivity. Through these effects, Bursicon signaling acts to promote metabolic adaptation to sugar intake and maintain glycemic control.

## Discussion

The global rise in obesity is fueling an increase in type-2 diabetes and other metabolic disorders, which are largely driven by resistance to insulin, the key hormone essential for governing glycemic levels and energy use and storage. Despite the importance of insulin resistance in pathogenesis, the molecular mechanisms underlying this phenomenon are not fully understood. In this study we performed a comprehensive *in-vivo* screen covering the secretome and receptome to identify hormonal mechanisms and pathways modulating sugar tolerance and cellular responses to insulin in a sugar-induced metabolic state characterized by obesity and resistance to insulin. Our findings provide important insight into how interorgan communication between the musculature, the nervous system, and adipose tissue influences sugar tolerance and insulin resistance, broadening our understanding of the intricate metabolic and endocrine networks involved in glucose homeostasis and shedding light on mechanisms relevant to diabetes pathogenesis.

We discovered that Rickets, the *Drosophila* ortholog of mammalian Leucine-rich repeat-containing G-protein coupled receptor 4 (Lgr4), is an important factor in metabolic regulation, promoting insulin production and adipose-tissue sensitivity to insulin. Lgr4 has been implicated in obesity-related metabolic dysfunction, and in humans, polymorphisms and gain-of-function mutations in this gene have been linked to obesity^45,46^. Lgr4 affects sugar tolerance and lipid metabolism and is generally believed to act through potentiation of Wnt/β-catenin signaling^47^. However, Lgr4 has also been suggested to mediate signaling through the cAMP-PKA pathway. Our findings suggest that the *Drosophila* homolog of Lgr4, Rk, regulates insulin production through a mechanism that involves the cAMP-PKA pathway, which is known to stimulate insulin secretion in mammalian β-cells^48^. Lgr4 is expressed in β-cells^49^, although its function in these cells remain unclear. Based on our findings, it will be interesting for future studies to explore whether Lgr4 promotes insulin secretion from mammalian β-cells via activation of the cAMP-PKA pathway. Together, our work provides evidence for the importance of Lgr signaling in metabolic regulation and suggests the intriguing possibility that modulation of Lgr4 signaling could potentiate insulin secretion and promote sensitivity to insulin in adipose tissue, improving glycemic control in high-sugar diet conditions that cause hyperglycemia.

Our results implicate the secreted lipocalin NLaz, homologous to human ApoD, as a possible factor in HSD-induced phenotypes and in their suppression by Bursicon signaling in adipose tissue. Previous reports have shown that NLaz affects longevity, stress resistance, and metabolism in *Drosophila*, and ApoD has been linked to obesity and insulin resistance, but the role of ApoD in diabetes and metabolic disorders remains to be clarified ^9,50,51^. Our work suggests that downregulation of NLaz downstream of Rk-mediated Bursicon signaling in the adipose tissue is necessary for the maintenance of glycemic control under conditions of high-sugar diet. These findings provide a molecular context for understanding the mechanism by which ApoD/NLaz and Rk/Lgr4 signaling regulate metabolic signaling in obese-like states and suggest that modulation of NLaz/ApoD function might provide a strategy for treatment of metabolic disorders.

Emerging evidence suggests that complex interactions between metabolism and the immune system are involved in metabolic homeostasis. Our findings shed new light on this interplay by revealing a previously uncharacterized interaction between NF-κB/Relish and insulin signaling. NF-κB/Relish mediates inflammatory and immune responses, and adipose-tissue inflammation is a central feature of obesity that contributes to insulin resistance. Accumulating evidence suggests that dysregulation of NF-κB signaling is linked to metabolic disorders, and studies in mammals have indicated that inhibiting NF-κB in relation to obesity-induced adipose inflammation could potentially provide metabolic benefits^52^. Our findings suggest that NF-κB/Relish inhibition may indeed improve insulin signaling and insulin sensitivity in adipocytes in obesity-like states, implying that NF-κB inhibition may alleviate adipose-tissue insulin resistance in obesity. The mechanism by which NF-κB/Relish-pathway modulation affects insulin resistance represents an interesting question for future investigations.

Maintaining metabolic homeostasis under nutritional stress requires a network of interorgan crosstalk to ensure coordinated adaptive responses of different organs and to effectively balance the uptake, use, and storage of energy. Given this network’s complexity, signaling routes that connect and coordinate the functions of organs to maintain metabolic homeostasis have remained difficult to elucidate. Our work here describes complex communication between muscle, neurons, and adipose tissue that is crucial for metabolic adaptations to a high-sugar diet. This supports an emerging paradigm in which metabolic control requires a coordinated effort by multiple organs that each sense different aspects of nutritional intake and metabolic state and relay information to other tissues to balance energy storage and mobilization. In both flies and mammals, fat-derived hormones (leptin or Unpaired 2) convey metabolic information and act via a neuronal relay to regulate insulin secretion^23,53^. Leptin also modulates the insulin sensitivity of skeletal muscles via central relays and thus mediates communication about the metabolic state of the fat tissue via neurons to both insulin-producing β-cells and the muscles to modulate their response to insulin. Our findings add a new axis to the model for regulating glucose homeostasis by demonstrating the existence of a muscle-derived signal that acts via a neuronal relay to modulate both insulin secretion and adipose tissue sensitivity to insulin. Our data suggest that, in flies, Gbb, a conserved BMP5/6/7/8 ortholog, is a sugar-regulated myokine that acts through its receptors (Tkv and Wit) on Bursicon-expressing neurons, which in turn drive both insulin production and adipose insulin sensitivity. All of the insulin- and glucose-homeostasis-regulating genes and pathways investigated in the present work have mammalian orthologs, which is not surprising given the evolutionary conservation of insulin signaling and the central importance of metabolic regulation. This suggests that these overall mechanisms and tissue-crosstalk routes might be conserved across species. The mammalian Gbb ortholog BMP7 has been shown to improve insulin signaling in insulin-resistant cells^54^, suggesting that this might be a conserved function of Gbb/BMP7. Although the function of BMP7 in energy metabolism is not well-characterized, it seems to act through leptin-independent mechanisms, making it of therapeutic interest in obesity, since the obese state is often characterized by leptin resistance^55^. Identifying the source of insulin-sensitivity-modulating BMP7 will be an interesting avenue of future research, as will examining whether the effects of BMP7 in this regard are mediated by downstream Lgr signaling.

In summary, this work unravels an intricate muscle-neuronal-adipose communication mechanism that involves BMP, Lgr, ApoD, and NF-κB components and pathways. This axis regulates glucose homeostasis under conditions that drive the pathological hallmarks of diabetes, including tissue resistance to insulin, by governing both insulin secretion and the insulin sensitivity of adipose tissue. Uncovering these mechanisms is not only fundamentally important but may also facilitate the development of targeted interventions for obesity and related metabolic disorders.

## Methods

### *Drosophila* husbandry and stocks

Flies were maintained on a standard lab diet (8.2% cornmeal, 6% sucrose, 3.4% baker’s yeast, and 0.8% agar, with 0.48% propionic acid and 0.16% methyl-4-hydroxybenzoate, based on a previous report^9^) on a 12/12-hour light cycle at 25 °C and 60% relative humidity. This diet, defined as “1x sugar”, is the basis for higher-sugar media: *e.g*., for “5x sugar” diet, sucrose was increased to 30%, with other ingredients left unchanged. Stocks obtained from the University of Iowa Bloomington *Drosophila* Stock Center (BDSC) include: *Burs^Z4410^*, likely a null allele due to mis-splicing^56^, #66432 (*w; arm-GFP; Burs^Z4410^/TM6B, Tb^+^*). *Arm-GFP* was removed, and the balancer was replaced with *TM6B, Tb*. *Burs-GAL4*, #40972; CaLexA system^43^, #66542, modified by adding *10xUAS-IVS-myr::tdTomato[su(Hw)attP8]* (#32223) as described in Malita *et al.*^22^*;Cg-GAL4*^57^, #7011; *daughterless (da)-GAL4*, #55850; *gbb-RNAi*, #34898; *Ilp2-GAL4*^58^, #37516; *Mhc-GAL4/TM3, Sb,* #55133, outcrossed five times to our standard lab genetic background to remove an unlinked homozygous-lethal allele and re-balanced over *TM6B, Hu Tb. UAS-PKA-R1^BDK^* ^59^, #35550; *R57C10-GAL4*^60^, *#*39171; *tGPH*^40^ *(alphaTub84B-GFP::PH(Grb1)*), #8164; *trans-Tango* system^36^, #77124; and *Tub-GAL80^TS^* ^61^, #7018, #7019. Stocks obtained from Vienna *Drosophila* Resource Center (VDRC) include *Burs-RNAi*, # 102204; *foxo-RNAi*, #107786; *NLaz-RNAi*, #101321; *Pburs-RNAi*, #102690; *rel-RNAi*, #108469; *rk-RNAi*, #105360; *tkv-RNAi,* #105834; and *wit-RNAi, #*103808. Other RNAi lines are listed in Table S1. *UAS-foxo* has been described previously^62^. *Rickets-GAL4*^63^ was a kind gift of Ben White (NIH). *UAS-DILP2HF*^64^ was a generous gift of Sangbin Park and Seung Kim (Stanford).

### High-sugar screen

A list of secretome and receptome genes was generated using the online resources GLAD^65^ (“Secreted Proteins” and “Receptors”) and MetazSecKB^66^ (“Highly likely secreted” and “Plasma Membrane” with probability ≥3). To expand the list of secreted proteins, we also included genes annotated as “Secreted” or associated with the GO terms “Extracellular region” (GO:0005576), “Extracellular space” (GO:0005615), or “Extracellular matrix” (GO:0031012) in FlyBase, UniProt, and Ensembl. These lists were merged to create one common list and cross-referenced with stock availability from Vienna *Drosophila* Resource Center (VDRC)^67^. One RNAi line was chosen for each gene, with lines from the KK collection preferred over those of the GD library. Additional lines from the University of Indiana Bloomington *Drosophila* Stock Center (BDSC) were included for genes from the GLAD Receptome list for which no VDRC RNAi stocks were available. The list of genes and RNAi lines is available as Supplemental Table S1.

Four males of each RNAi line were crossed to six *da-GAL4* virgin females in vials containing 1x- and 5x-sugar medium, and flies were allowed to seed the vials with eggs for 24 hours at 25 °C. Flies were transferred to new vials at least twice for additional egg-lays. Adults were removed, and the vials were incubated at 25 degrees. The formation of prepupae or pupae (marked by visible cuticle darkening, Bainbridge and Bownes^68^ stage 9) was recorded once each day. Several vials of each cross were scored, and the mean time until 50% pupariation was calculated for replicates that had a minimum of three pupae using a custom written MATLAB script (the MathWorks, Natick, MA). Other defects such as larval arrest were also recorded. Genotypes that exhibited a phenotype on 5x sugar but not on 1x sugar – those that exhibited a sugar-dose-specific phenotype – were considered to be of interest for follow-up.

### Other pupariation assays

Crosses were set up in egg-laying chambers sealed with a 60-mm Petri dish containing apple-juice agar (1 L water, 340 mL apple juice, 30 g agar, 34 g glucose, 20 mL 10% Tegosept in ethanol). Animals were allowed to lay eggs for four hours at 25 °C, and the plates were incubated for 24 hours at 25 °C. Hatched larvae were transferred to vials containing 1x or 5x medium, with ∼30 larvae per vial, and vials were incubated at 25 °C or 29 °C. Pupae (completion of spiracle eversion and complete immobilization of animals) were counted every 4-8 hours. At least 5 vials were scored for each genotype.

### ELISA of circulating tagged DILP2

A stock of *Ilp2-GAL4, UAS-Ilp2(HA,FLAG)*^64^ was created, and this was crossed to *w*^1118^ or to *UAS-rk-RNAi*. Four-hour egg-lays were made, and larvae were transferred 24 hours later to vials containing 1x-sugar medium (∼30 animals per vial). At 96 hours after the midpoint of the egg-laying period, mid-third-instar larvae were recovered from the medium, washed with MilliQ water twice to remove any particles of the medium, and dried on a piece of paper towel; hemolymph was collected by cutting the larval cuticle with iridotomy scissors and collecting the exudate on an ice-cold shallow-welled glass slide. Samples were heat-treated at 60 °C for 5 minutes to denature phenoloxidase (melanization enzyme), centrifuged to remove any hemocytes, debris, or aggregates, and used in a sandwich HA/FLAG ELISA. Anti-FLAG (Sigma-Aldrich #F1804, 5 μg/mL in 200-mM NaHCO_3_ buffer, pH 9.4) was adsorbed onto F8 MaxiSorp Nunc-Immuno modules (ThermoScientific #468667) overnight at 4 °C. Modules were washed twice with phosphate-buffered saline (PBS) + 0.1% Triton X-100 (PBST) and then were blocked for 2 hours at room temperature with PBST + 4% non-fat dry milk. Modules were washed three times in PBST, and 1 μL of hemolymph or synthetic HA::spacer::FLAG peptide standard (DYKDDDDKGGGGSYPYDVPDYA) was diluted into 50 μL PBST + 25 ng/mL mouse anti-HA peroxidase (Roche, #12013819001) + 1% non-fat dry milk, added to the wells, and incubated overnight at 4 °C. The solution was removed, and the modules were washed six times, five minutes each, with PBST. One hundred microliters of One-step Ultra TMB ELISA substrate (Thermo Scientific #34028) was added to each well, and the modules were incubated for 15 minutes at room temperature to permit color development. The reaction was terminated by the addition of 100 μL 2 M sulfuric acid, and the absorbance at 450 nm was measured using an EnSight plate reader (PerkinElmer).

### Metabolite analyses

Triacylglycerides (TAG) and glucose levels were measured using standard colorimetric methods^69,70^. Four-hour egg lays were performed, and at 24 hours after egg laying (AEL), larvae were transferred to vials of 1x or 5x-sugar medium, ∼30 per vial. In chronic-feeding experiments, vials were incubated at 25 °C until 96 hours after egg laying for 1x sugar or 144 hours AEL for 5x sugar, to equalize developmental progression; for short-term 5x-sugar feeding, animals were raised on 1x diet until 90 hours AEL and then transferred to fresh 1x medium or to 5x-sugar diet for ten hours (until 100 hours AEL). Several samples each containing 4 larvae were collected in 260 μl PBST (PBS+0.05% Tween-20, Sigma #1379) and homogenized using 5-mm steel beads (Qiagen #69989) in a bead mill (Qiagen TissueLyser LT). For TAG measurements, acylglyceride ester bonds were cleaved using Triglyceride Reagent (Sigma, #T2449) to liberate glycerol, and the concentration of this product was measured using Free Glycerol Reagent (Sigma, #F6428). Glucose was measured in a colorimetric assay (Sigma #GAGO20). For hemolymph assays, hemolymph was collected by cutting cuticle with micro scissors and collecting the exudate with a pipette? Hemolymph was 10-fold diluted with PBS and heat-treated at 70 °C for 5 minutes to prevent melanization and centrifuged to remove pelleted aggregates. Hemolymph glucose concentrations were determined using the same kit as above. In TAG and glucose assays, the resulting absorbance at 540 nm was measured in a 384-well plate using an Ensight multimode plate reader (PerkinElmer). Protein concentrations were determined using a bicinchoninic acid assay (Sigma, #BCA1), and the resulting absorbance was read at 562 nm. Absorbances were converted to concentrations using standard curves for glucose, glycerol, and protein. TAG and glucose levels are reported as the ratio of those metabolites to protein to normalize for slight differences in larval size.

### Transgene construction

*Transgenes for inducible CRISPR-mediated knockout:* UAS-driven guide-RNA constructs targeting two sites in each of *rickets*, *Burs*, and *Pburs* were created in the vector pCFD6^71^, obtained from AddGene (#73915; https://addgene.org/). For each gene, gRNA target sequences were identified using the E-CRISP algorithm^72^ (https://e-crisp.org/E-CRISP/). A portion of pCFD6 was amplified by standard PCR using pairs of oligos, given in Supplementary Table S2, in which these gene-specific target sequences were inserted (underlined). Each resulting fragment was inserted into BbsI-digested pCFD6 using GeneArt Gibson Assembly HiFi Master Mix (ThermoFisher, #A46628). Clones were sequenced, and correctly assembled clones were midi-prepped using Qiagen kits and integrated into the genome at the *attP2* third-chromosome site by BestGene, Inc (Chino Hills, CA).

*Creation of endogenous FLAG::Burs::HA knock-in allele:* We designed a CRISPR-Cas9-mediated knock-in construct that adds a FLAG tag to the N-terminus and an HA tag to the C-terminus of mature secreted Burs. The FLAG tag sequence was inserted after the putative signal sequence^31,73^, and the HA tag sequence was inserted right before the stop codon. Three silent mutations were introduced into the gRNA target sites within the coding sequence to prevent recutting of the insert. The *FLAG::Burs::HA* construct was flanked by 800-base-pair homology arms. The resulting knock-in sequence is given in Supplemental Table S2. This sequence was synthesized by ThermoFisher, transformed into competent cells, and midi-prepped for microinjection. We designed three pairs of primers that contains sense or antisense gRNA sequences (see primer sequences in Supplementary Table S2) and cloned into plasmid pCFD3^74^. Clones were sequenced, and one correct clone of each gRNA construct was midi-prepped. The knock-in construct and the three gRNA constructs were mixed 3:1:1:1 at a final total concentration of 500 ng/µl, and this plasmid solution was injected into *nos-Cas9* embryos in-house. Successful knock-in fly lines were identified by PCR confirming the presence of the FLAG-tag sequence.

### Quantitative RT-PCR

For measurement of transcript levels, six samples each containing 5 whole larvae or 5 larval dissected tissues (central neuronal systems or body carcasses) were disrupted in 350 μl lysis buffer from the NucleoSpin RNA kit (Macherey-Nagel, #740955) containing 1% beta-mercaptoethanol, using a Qiagen TissueLyser LT bead mill and 5-mm stainless steel beads (Qiagen, #69989). RNA was isolated from the samples using the NucleoSpin RNA kit, and cDNA was produced using the High-Capacity cDNA Synthesis Kit (Applied Biosystems, #4368814). Real-time PCR was performed using RealQ Plus 2× Master Mix Green (Ampliqon, #A324402) on a QuantStudio 5 machine (Applied Biosystems). Expression was calculated using the delta-delta-Ct method, with ribosomal-protein gene *Rp49* as control. The oligos are listed in Supplemental Table S2.

### Western blotting

Larvae were raised for 90 hours AEL on 1x diet with further transfer either onto 1x or 5x diet for 10 hours, until 100 hours AEL at collecting, and samples of four larvae each were homogenized in 100 μl 2× SDS sample buffer (Bio-Rad #1610737) containing 5% (355 mM) 2-mercaptoethanol using a Qiagen bead mill (TissueLyser LT) and 5-mm steel beads. Homogenates were denatured at 95 °C for 5 minutes, and insoluble material was pelleted by five minutes’ centrifugation at maximum speed. Samples were loaded into a precast 4–20% polyacrylamide gradient gel (Bio-Rad) and electrophoresed. Separated proteins were transferred to polyvinylidene difluoride membrane (PVDF, Millipore) using a Trans-Blot Turbo Transfer Pack (Bio-Rad, #1704158) on a Trans-Blot Turbo setup (Bio-Rad). The membranes were incubated in Odyssey blocking buffer (LI-COR, #927-40100) for one hour at room temperature. Blocking solution was poured away, and the blots were incubated with rabbit anti-phospho-Akt (Cell Signaling Technology, #4054; 1:1000) and mouse anti-alpha-Tubulin (Sigma #T9026, diluted 1:4000) in Odyssey blocking buffer + 0.2% Tween-20. The membranes were rinsed and washed for three times for 15 minutes each with PBST (PBS supplemented with 0.1% Tween 20) and incubated for 45 minutes with IRDye 800CW anti-rabbit (LI-COR #925-32210) and IRDye 680RD anti-mouse (LI-COR #925-68070) secondary antibodies, diluted 1:10,000 in Odyssey blocking buffer + 0.2% Tween-20. Membranes were rinsed and washed for three times for 15 minutes each with PBST buffer, avoiding light, and imaged using an Odyssey Fc imaging system (LI-COR).

### Immunostaining

Larvae were raised at 25 °C until 90 hours after egg-laying on 1x diet with further transfer either onto 1x or 5x diet and kept for further 10 hours. Larvae were extracted from the medium and rinsed with PBS. For *ex-vivo* insulin-stimulation assays, fat-body tissue was dissected in PBS on ice, rinsed twice with cold PBS and transferred into a shallow glass dish containing Schneider’s medium. Either pure Schneider’s (“mock”) or Schneider’s containing human insulin (Sigma-Aldrich #I9278, final concentration, 0.5 μM) was added, and tissues were incubated for 20 minutes at room temperature before fixation and further processing as below. For other experiments, tissues were dissected in cold PBS. Tissues were transferred to 4% paraformaldehyde (PFA) in PBS and incubated at room temperature, with agitation, for one hour. After PFA removal, tissues were rinsed and washed three times, for 15 minutes each time, with PBSTx (PBS+0.1% Triton X-100). Tissues were blocked for 30 minutes at room temperature in PBSTx with 5% normal goat serum (Sigma). Primary antibodies were diluted as detailed below in blocking buffer, and tissues were incubated in this solution overnight at 4 °C with agitation. Tissues were quickly rinsed with PBSTx and washed three times for 20 minutes each with PBSTx. Secondary antibodies were diluted 1:500 in PBSTx, and tissues were incubated in this solution overnight at 4 °C with agitation. The secondary staining solution was removed, and tissues were rinsed once quickly with PBSTx and washed three times for 20 minutes each with PBSTx. If phalloidin was included in the experiment, the first wash contained Alexa Fluor 647-conjugated phalloidin, diluted 1:1000 from the manufacturer’s recommended stock concentration. The last wash contained DAPI (Sigma, #D9542), diluted 1:500 in PBSTx. Tissues were mounted in ProLong Glass anti-fade mountant (Invitrogen, #P36984), using a 0.12-mm-thick spacer (Grace Bio-Lab, #654006) and glass cover slip, on poly-lysine-coated slides (VWR, #631-0107) that had been further coated with poly-L-lysine (Sigma, #P8920). Slides were allowed to cure for at least 24 hours at 4 degrees. Tissues were imaged using a Zeiss LSM 900 confocal microscope using 20x and 40x objectives. Z-stacks were captured with a one-micron Z-step. Samples that were to be compared against one another were prepared at the same time, using the same reagent mixes, and were imaged using the same settings.

The following primary antibodies were used: rabbit anti-Foxo, 1:500, kindly provided by Pierre Léopold (Institut Curie)^75^; mouse monoclonal anti-GFP (clone 3E6, ThermoFisher #A11120), 1:500: rat monoclonal anti-HA (clone 3F6, Roche #11867423001), 1:1000: rabbit anti-Ilp2^76^, kind gift of Ernst Hafen and Michael Pankratz (University of Bonn), 1:1000; mouse anti-Ilp3^77^, kind gift of Jan Veenstra (Bordeaux), 1:1000; rat anti-Ilp5^78^, 1:500; mouse monoclonal anti-Relish-C, 1:50, from DSHB (21F3-c); and against tdTomato, rat anti-mCherry (ThermoFisher, no. M11217), 1:1000. The following secondary antibodies were used, all diluted 1:500: Alexa Fluor 488-conjugated goat anti-mouse (ThermoFisher, #A32723); Alexa Fluor 555-conjugated goat anti-rabbit (ThermoFisher, #A32732); Alexa Fluor 555-conjugated goat anti-rat (ThermoFisher, #A21434); and Alexa Fluor 647-conjugated goat anti-rat (ThermoFisher, #A21247)

### Image analysis

Image analysis was performed using FIJI/ImageJ^79^, version 1.53t. For quantification of Ilp staining in the IPCs: Z-stacks were projected using the “sum” method. Each IPC cluster was manually segmented using the freehand drawing tool, and the fluorescence values within this region of interest were summed. The region of interest was moved to an adjacent non-stained region, and the fluorescence within this area was summed as a background value. This background was subtracted from the value obtained for the IPCs. For measuring CaLexA or Bursicon levels: Z-stacks were projected using the “sum” method. Each cell of interest was located using the tdTomato channel and highlighted as a region of interest. The fluorescence of the GFP or anti-HA (Bursicon) channel within this region was summed. The region of interest was offset to a nearby unstained region, and the background intensity was summed and subtracted from the value obtained for the stained region. For quantification of tGPH localization, locally flat regions of fat-body tissue were identified in each Z-stack. For each of several pairs of adjoining cells in this region, using the phalloidin and DAPI channels, a Z section was selected at the depth of the nuclei, where the cell membranes are completely perpendicular to the section. A line (with a width of 15 pixels, to reduce noise) was drawn from one nucleus to the other, placing the midpoint of the line at the membrane, and the fluorescence intensity in the GFP channel at each pixel along this line was recorded. The positions along the line were normalized to a length of 1.0. The fluorescence values from the middle 10% of the line (that is, 0.45 ≤ *x* ≤ 0.55) were averaged to create a “center” intensity, and the fluorescence values within a further 10% on each side of this interval (that is, from 0.35 ≤ *x* < 0.45 and from 0.55 < *x* ≤ 0.65) were averaged to produce a “surround” value. The center:surround ratio, reflecting the level of membrane enrichment, is reported in the figures.

### Statistical analyses

Statistical calculations were performed using the *Prism* software package (GraphPad). All data sets were evaluated for normality before other tests were performed. The statistical tests for significance that were employed are listed in each figure are given in the respective legend.

## Data Availability

All data created for this work are available in this manuscript, its figures, or its supplemental files. Raw images may be obtained on reasonable request to the corresponding author.

## Code Availability

The custom MATLAB script used for sleep analysis in this study is available in Supplementary Table S3.

## Acknowledgements

*Rickets-GAL4* (*rickets^Pan^-GAL4*)^63^ was a kind gift of Ben White (NIH). Rabbit anti-Ilp2^76^ was a gift of Ernst Hafen and Michael Pankratz (University of Bonn). Mouse anti-Ilp3^77^ was a gift of Jan Veenstra (University of Bordeaux). Rabbit anti-Foxo was a gift of Pierre Léopold (Institut Curie). Plasmids pCFD3 and pCFD6 were obtained from AddGene. Fly stocks were also obtained from University of Indiana Bloomington *Drosophila* Stock Center and Vienna *Drosophila* Resource Center. Anti-Relish was obtained from University of Iowa Developmental Studies Hybridoma Bank. This work was supported by the Danish Council for Independent Research Natural Sciences grant 8021-00055B to K.R. and Novo Nordisk Foundation grant NNF21OC0070402 to KR. A.J.F was supported by a Novo STAR co-financed industrial PhD fellowship from Novo Nordisk Foundation and Innovation Foundation. T.K. and K.V.H. were supported by funding from the Danish Council for Independent Research Natural Sciences (grant 9064-00009B) to K.V.H.. The Zeiss LSM 900 confocal microscope and the PerkinElmer EnSight plate reader were purchased with equipment grants from the Carlsberg Foundation (CF19-0353 and CF17-0615) to K.R. et al.

## Author contributions

A.F.J., M.J.T., J.L.H., and K.R. conceived and designed the study. O.K., A.F.J., M.J.T., T.K., S.N., M.L., J.L.H., and K.R. designed, performed, and analyzed experiments. J.H. and D.M. contributed to the design of the list of genes. M.J.T. and K.R. wrote the manuscript, and O.K., A.F.J., T.K., S.N., J.H., M.L., K.V.H., and J.L.H. reviewed and edited the manuscript.

## Extended data

**Extended data, Figure 1.**
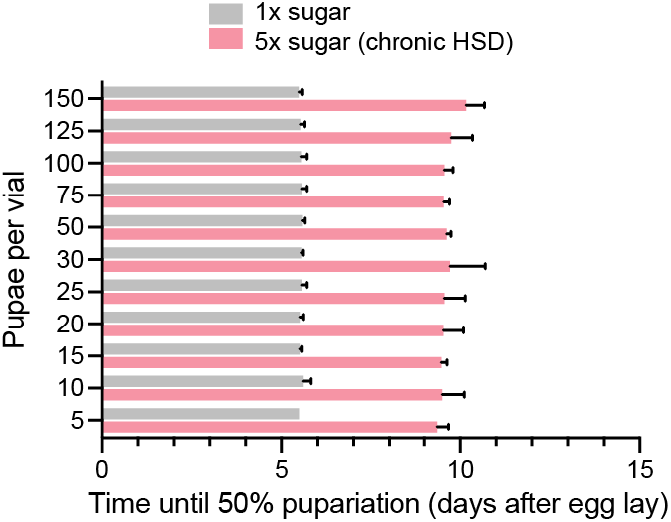
Pupation timing for animals raised on normal or high-sugar diet, at a range of population densities. Statistics: Error bars represent mean and SEM.

**Extended data, Figure 2.**
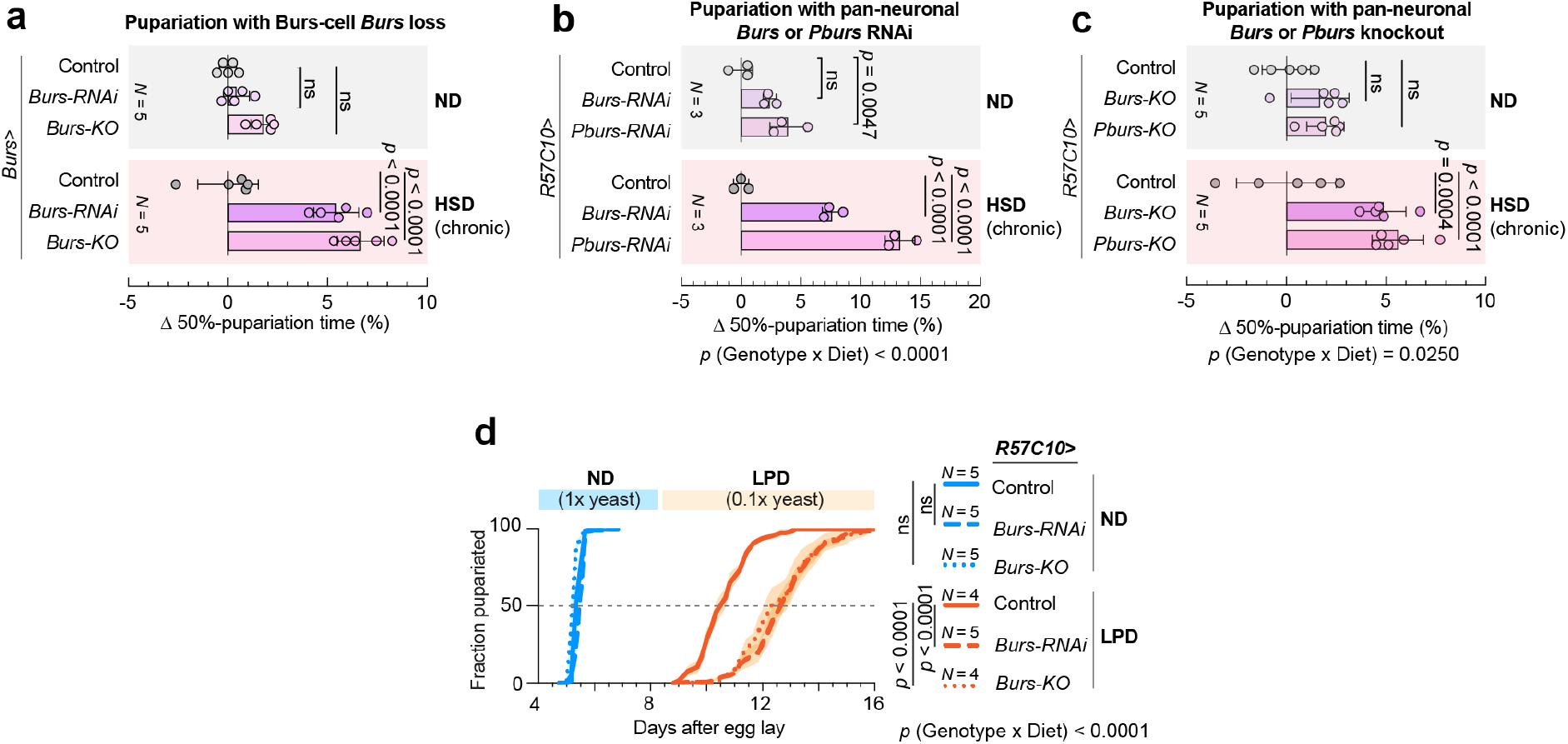
(A) Related to Figure 2B. Difference in the time until 50% pupariation between controls and animals lacking *Burs* expression in the *Burs*-expressing cells, raised on normal or high-sugar diet. (B) Difference in the time until 50% pupariation between controls and animals expressing pan-neuronal knockdown of *Burs* or *Pburs*, raised on normal or high-sugar diet. (C) Difference in the time until 50% pupariation between controls and animals expressing pan-neuronal somatic deletion of *Burs* or *Pburs*, raised on normal or high-sugar diet. (D) Pupation timing on normal and low-yeast diets for animals with pan-neuronal knockdown or knockout of *Burs*. Statistics: Error bars represent mean and SEM. ns, not significant (p>0.05). A, B, C, D, one-way ANOVAs with Tukey’s correction between 50%-pupariation times for multiple comparisons and two-way ANOVA for interaction.

**Extended data, Figure 3.**
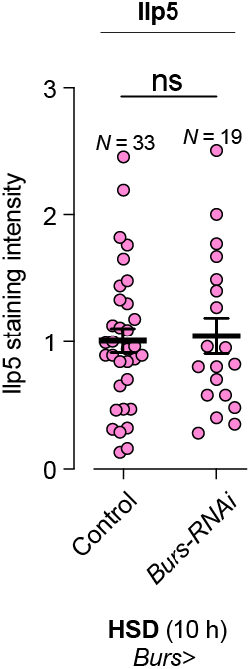
Quantification of anti-Ilp5 staining intensity in multiple samples from animals with Burs^+^ cell *Burs* loss. Statistics: Error bars represent mean and SEM. ns, not significant (p>0.05). Two-sided unpaired t-test.

**Extended data, Figure 4.**
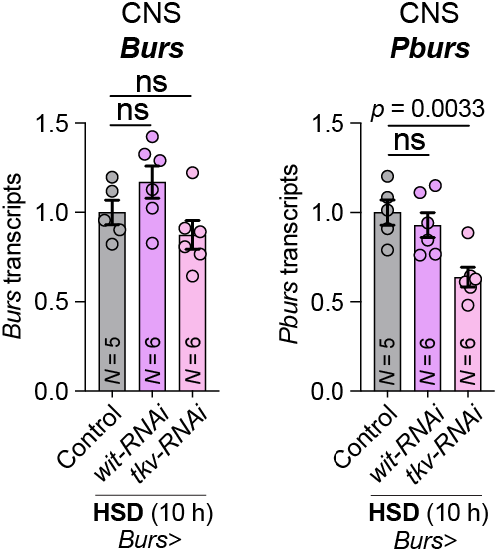
Transcript levels of *Burs* (left) and *Pburs* (right) measured by qPCR in dissected CNS samples from controls and animals expressing RNAi against *wit* or *tkv* in the *Burs*-expressing cells, fed high-sugar diet for 10 hours. Statistics: Error bars represent mean and SEM. ns, not significant (p>0.05). One-way ANOVAs with Dunnett’s correction.

**Extended data, Figure 5.**
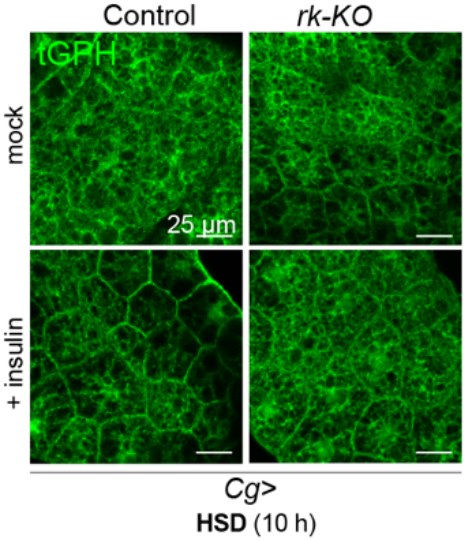
Representative images of tGPH insulin-indicator responses to exogenously applied human insulin in fat-body explants from controls and animals expressing CRISPR-mediated deletion of *rickets* in the fat body, exposed to high-sugar diet for 10 hours before dissection. Scale bars, 25 microns.

