## Supplementary Table 2 for "A neuronal relay mediates muscle-adipose communication that drives systemic metabolic adaptation to high-sugar diets"

**Supplementary Table S2.**

Oligos for gRNA constructs

*Burs-KO-F:* CGGCCCGGGTTCGATTCCCGGCCGATGCAGATTCCCTCGTTCGCCTGCGGTTTCAGAGCTATGCTGGAAAC

*Burs-KO-R:* ATTTTAACTTGCTATTTCTAGCTCTAAAACCAGGAAATTGCCGGCTATTCTGCACCAGCCGGGAATCGAACC

*Pburs-KO-F*: CGGCCCGGGTTCGATTCCCGGCCGATGCAGAAGCATGGATATACGTCTTGTTTCAGAGCTATGCTGGAAAC

*Pburs-KO-R:* ATTTTAACTTGCTATTTCTAGCTCTAAAACCCTTGATCAAGTGGATCTCCTGCACCAGCCGGGAATCGAACC

*rk-KO-F:* CGGCCCGGGTTCGATTCCCGGCCGATGCAGTTGCTACCTGAAGATGTACGTTTCAGAGCTATGCTGGAAAC

*rk-KO-R:* ATTTTAACTTGCTATTTCTAGCTCTAAAACTTCACCTGAACAAGAGGCTCTGCACCAGCCGGGAATCGAACC

FLAG::Bursicon::HA knock-in sequence

Homology1 UTR Signal sequence FLAG tag Burs intron Burs intron Burs HA tag UTR homology2

TACGAGGAGAATCCTGACTTGGTTCAGGCAATTTTCGAAATGATTACCGAAAACATTATGGAATTAGCGTATAAGTGAGTTATTTTTCATAATTTGATCATCTGAGAGTGTGGTTTGATTGTTTATTGAACACTTAACCTTGTCGCTTTTCAGCTTCCAGAGTTGGTCGGGACACGGCCGAGGTGATCAATGACATGGCGGAGTACTTCACCACCCGGGAGCAACAGCAGTTGCTCAAGGAATTCTACTACAGCAATCACCAGTTCTTCGGTGAATCGGAAGAGCTGCTCTCCAAGTCCCTGGTCACTGTGGAAAAAAACCTTAACTGGGCGGAAACCCATTTGGAAGGCTTAGTAAAATATCTGGCGGACAGAAATGGTAGCTCCTGCCTGGGAGCCACTTCCTTGGTGGTGGTTCTGCTTTTAGCAGTGGGCAGCATGTTGGCCTGGCCCTGATTAGCCAATAAGTTGTGAGGGAAATAAAGAGATTGCCGTGGTCGTGCGCCGCGGGAGAAATGCCCTTTTCGTGTACGCAATTACTAGGCAAAAAGGTTTGGACGGCCCTTGCTTTATTAGGCCATAATTTACGCTGTCGAGCGGCCATTAGAGCGGCATCCGCTGGCCAAGTTGGTCTATAAAAGCCCGGCGGCTGCAAGTGGCGTTTTCCATTCCACGTGAAAGGACACTCGCAGTCGGGCCGACGAG**ATG**CTGCGCCACCTGCTCCGCCACGAGAACAACAAGGTCTTCGTCCTGATCCTGCTCTACTGCGTCCTGGTCAGCATTCTGAAACTCTGCACGGCAGACTACAAAGACGATGACGACAAGCAGCCGGATAGCTCTGTGGCCGCCACGGATAATGGTATTTGATAAGGCATTTGATAATATCATATTATTCGCTCTAATGGATTCCATTTACCATCCAGATATTACGCATCTTGGCGACGATTGTCAGGTGACGCCCGTCATCCATGTGCTCCAGTATCCTGGATGTGTGCCCAAGCCGATTCCCTCGTTCGCCTGCGTGGGTCGCTGTGCCAGTTATATCCAGGTGGGTGTTGATATACTTGAAATATAAATTATAATTTGATATAATCTGATGACTTTAGGTTTCGGGCAGTAAGATCTGGCAAATGGAGCGTTCCTGCATGTGCTGCCAGGAGTCTGGTGAGCGGGAGGCAGCCGTCTCGCTATTCTGTCCCAAAGTGAAGCCCGGCGAGCGTAAATTCAAGAAGGTCCTGACCAAGGCGCCATT[G🡪A]GAGTGCATGTGTCGGCCATGCACTTCCATTGAGGAGTCTGGCATCATACCACA[G🡪A]GAAATTGCCGGCTATTCGGACGAGGGTCCACTCAACAATCACTTCCG[G🡪C]CGCATTGCTCTGCAATACCCATACGATGTTCCAGATTACGCT**TAG**ATTCCCCCATCAGTTTAGCACTCATACCCATGCCCATTTGCTCATTAAAATAAATTAGAGTTGCCGTTTCTGCTTGTACACAGTGAAAAATGTGCCTCTGAAAGCTCATGCCAGTGAAAGGGAAACATCAGCAGATATATGATTATTTTAAATTGTAATTTATTTCCCAACTATCTCATGTCATTGCTACTAGTTCACTAAATGGTGTGTGTCCTGCTTCACAGGGTATGTATGTAGATGCGTAACTCCAAAGAAAACAAAGATAATCTTTACACTGCAACTGGATCTAACAATAGGAGTTTAACTACGGCATAATGCATATGACTCGCGCCGCCTACAAGTACTCGACGATATCGCATTTGATGGATATATTCGGACAGCTCTGGCCCACGGCACTGCGGCCCCGATAGTTGGGACTCTTCTCGGTGAGGAGTTTCCGGAAGATGGGCAAAATTGCCGTCTCGCCCTCGAGCAGTTCCGTCTCGCTGCTGGCGCCGTAGATGCTGCTATCCTGCTTGCCGCGCTGCCTGCTCCGCTGCTTATCGGGCGCCACATTGGGCGAACTGTGGTGGTGGGCGGCGTGCTGGATGTTAGAGCTCCGGCTGACCTTACGCGGCTGGACGCACGGCTGACTGTTGGGCGGCGGATCTCGTTCCCGTTCGCTGTCCGTCTCGCTCTGGCAAACGGTGCCCTTTTGCGGTTTTGGCCGATGTTTGTTGCGGTTCTTCTCGGAGACGGACCGCTGGATGGGCGAGGTGGTGTTCGGGTCATAGGTCATCCCGTGGCCGCCATGATGT

Oligos for CRISPR knock-in

Burs-gRNA1-F: GTCGTCCACTCAACAATCACTTC

Burs-gRNA1-R: AAACGAAGTGATTGTTGAGTGGA

Burs-gRNA2_F: GTCGGAGTCTGGCATCATACCAC

Burs-gRNA2_R: AAACGTGGTATGATGCCAGACTC

Burs-gRNA3_F: GTCGGTCCTGACCAAGGCGCCAT

Burs-gRNA3_R: AAACATGGCGCCTTGGTCAGGAC

Oligos for qPCR

4EBP-F: TGCCCATGATCACCAGGAAG

4EBP-R: TCGTAGATAAGTTTGGTGCCTCC

Burs-F: CATCCATGTGCTCCAGTATCC

Burs-R: GGCTTCACTTTGGGACAGAA

gbb-F: GCGAGTGCAATTTCCCGCTCAATG

gbb-R: CAGGTTCACATTCTCGTCGTTCAGG

Ilp2-F: CTCAACGAGGTGCTGAGTATG

Ilp2-R: GAGTTATCCTCCTCCTCGAACT

Ilp3-F: AGAGAACTTTGGACCCCGTGAA

Ilp3-R: TGAACCGAACTATCACTCAACAGTCT

Ilp5-F: ATGGACATGCTGAGGGTTG

Ilp5-R: GTGGTGAGATTCGGAGCTATC

NLaz-F: ACGCCAACTACAGTCTCATAGA

NLaz-R: CGAGGGTTGTCCGGTGAATC

Pburs-F: AGGATTGTGCAACAGTCAGG

Pburs-R: AGCAATGGGTTAGAGTGATGA

Rp49-F: AGTATCTGATGCCCAACATCG

Rp49-R: CAATCTCCTTGCGCTTCTTG
