## Supplementary Table 3 for "A neuronal relay mediates muscle-adipose communication that drives systemic metabolic adaptation to high-sugar diets"

%% Calc mean Dev time for 5x and 1x

index=unique(Matlab51Subset.Symbol);

for ii =1:length(index);

Average5 (ii)= nanmean([Matlab51Subset.Dev5A(Matlab51Subset.Symbol==index(ii)); Matlab51Subset.Dev5B(Matlab51Subset.Symbol==index(ii)); Matlab51Subset.Dev5C(Matlab51Subset.Symbol==index(ii))]);

end

for ii =1:length(index);

Average1 (ii)= nanmean([Matlab51Subset.Dev1A(Matlab51Subset.Symbol==index(ii)); Matlab51Subset.Dev1B(Matlab51Subset.Symbol==index(ii)); Matlab51Subset.Dev1C(Matlab51Subset.Symbol==index(ii))]);

end

%% average logical (arrest)

for ii =1:length(index);

Arrest5(ii)={[Matlab51Subset.Arrest5A(Matlab51Subset.Symbol==index(ii)), Matlab51Subset.Arrest5B(Matlab51Subset.Symbol==index(ii)), Matlab51Subset.Arrest5C(Matlab51Subset.Symbol==index(ii))]};

end

for ii =1:length(index);

Arrest1(ii)={[Matlab51Subset.Arrest1A(Matlab51Subset.Symbol==index(ii)), Matlab51Subset.Arrest1B(Matlab51Subset.Symbol==index(ii)), Matlab51Subset.Arrest1C(Matlab51Subset.Symbol==index(ii))]};

end

%% combine data into dataset

averagedata=dataset(index,Average5',Arrest5', Average1', Arrest1');

averagedata.Properties.VarNames{1} = 'Symbol';

averagedata.Properties.VarNames{2} = 'MeanDev5';

averagedata.Properties.VarNames{3} = 'Arrest5';

averagedata.Properties.VarNames{4} = 'MeanDev1';

averagedata.Properties.VarNames{5} = 'Arrest1';

export(averagedata,'XLSFile','\\FSDKHQ001\Users405$\AEFJ\FollowMe\data\test.xlsx')
